# PCIF1 loss licenses antitumour immunity via cholesterol biosynthesis

**DOI:** 10.64898/2026.07.11.737975

**Authors:** Kaixiu Li, Tianjian Lu, Zheyu Ding, Mengyao Wang, Séverine Roy, Yulan Deng, Naima Benannoune, Yuqing Wang, Elodie Edmond, Jean-Yves Scoazec, Caroline Robert, Lunxu Liu, Shensi Shen

**Affiliations:** Department of Thoracic Surgery and Institute of Thoracic Oncology, Frontiers Science Center for Disease-related Molecular Network, West China Hospital, Sichuan University, Chengdu, China; Oncology Department, Dermatology Unit, Gustave Roussy, Villejuif, France; Department of Pathology, Gustave Roussy, Villejuif, France; Paris-Saclay University, Orsay, France; INSERM U981, Villejuif, France

## Abstract

Cholesterol biosynthesis is classically viewed as a tumour supportive metabolic programme, yet cholesterol-associated metabolic states can paradoxically coincide with improved responses to immune checkpoint blockade (ICB)^1^, highlighting an unresolved context-dependent relationship between cholesterol homeostasis and antitumour immunity. Here we identify PCIF1, a cap-specific RNA methyltransferase^2^, as a tumour-intrinsic brake on the inflammatory potential of cholesterol biosynthesis. PCIF1 depletion had limited effects on tumour cell proliferation *in vitro*, but suppressed tumour growth in immunocompetent, not immunodeficient, hosts and sensitized immune-refractory syngeneic tumours to ICB. Immune profiling by single-cell RNA sequencing, flow cytometry and multiplex immunofluorescence revealed increased CD8+ T cell infiltration in PCIF1-deficient tumours, and CD8+ T cell depletion restored tumour growth. Mechanistically, PCIF1 loss selectively enhanced translational engagement of SCAP, activated SREBP2-dependent cholesterol biosynthesis and promoted intracellular cholesterol accumulation. Unexpectedly, activation of this cholesterol programme induced a tumour cell inflammatory cytokine state enriched for T cell-recruiting chemokines, including CXCL10. Pharmacological perturbation of the SCAP-cholesterol axis attenuated this inflammatory programme, placing cholesterol sensing upstream of PCIF1 loss-induced tumour cell inflammatory activation. In ICB-treated patient cohorts, high PCIF1 expression in pre- and on-treatment samples was associated with poor clinical response, and across tumour types, PCIF1 expression inversely correlated with cytolytic immune signatures. These findings reveal PCIF1-mediated SCAP translational control as a tumour cell checkpoint that constrains the antitumour inflammatory activity of a canonical growth-associated cholesterol programme and shapes checkpoint therapy responsiveness.

## Introduction

Durable benefit from immune checkpoint blockade (ICB) is limited by tumour beds that are not permissive to cytotoxic T cell entry and function^3^. Tumour cell metabolic state can shape this barrier^4^, but its relationship to antitumour immunity is not always linear. Cholesterol biosynthesis is essential for membrane biogenesis and tumour fitness^5^, and cholesterol-linked pathways can promote malignancy in glioblastoma, bladder cancer and other settings^6–8^.

Conversely, cholesterol accumulation can enhance CD8□ T-cell cytotoxicity, NK-cell function and dendritic-cell immunogenicity^9–11^. Clinical observations linking cholesterol-associated metabolic states to improved ICB response further suggest that cholesterol biosynthesis can intersect with immune-responsive tumour states^1,12^. What links this lipid homeostatic programme to tumour inflammation remains poorly understood. Here we show that tumour cell PCIF1 restrains this connection. PCIF1 is a cap-specific mRNA methyltransferase that deposits N^6,2^-O-dimethyladenosine (m□Am) at the 5′ end of transcripts^2,13^, a position that places it close to transcript fate and translational control^14^. PCIF1 has been linked to viral immune evasion and attenuation of interferon responses^13,15^, but whether it acts within tumour cells to regulate antitumour immunity is unknown. Our studies identify a PCIF1-regulated translational layer that gates the inflammatory output of cholesterol biosynthesis and shapes cytotoxic T cell permissive tumour states and immune checkpoint responsiveness.

### PCIF1 loss promotes immune-dependent tumour control

We first asked whether tumour cell PCIF1 contributes to immune evasion in vivo. To this end, we stably depleted Pcif1 using lentivirus-based shRNA in three syngeneic murine tumour models that span distinct immunotherapy response states: the ICB-sensitive *Braf^V600E^/Pten^-/-^*melanoma (BP)^16^, the less responsive *Braf^V600E^/Pten^-/-^/Cdkn2^-/^*^-^ melanoma model (Yumm1.7)^17^ and the immune-refractory *Kras^G12C^*-mutant Lewis lung carcinoma model (LLC1)^18^ (**Extended Data Fig 1a**). This panel allowed us to test whether the effect of PCIF1 loss was restricted to a single tumour lineage or instead reflected a broader tumour cell intrinsic mechanism operating across different immune contexts. Pcif1 depletion did not affect the *in vitro* proliferation across these models (**Extended Data Fig 1b**). We next implanted control and Pcif1-depleted tumour cells into immunocompetent syngeneic *C57BL/6J* mice. In this setting, Pcif1 loss delayed tumour outgrowth across all three models compared with non-targting control cells (shNTC) (**Fig 1a-c**). The growth restraint was accompanied by prolonged survival of tumour-bearing mice (**Fig 1d-f**), indicating that tumour cell PCIF1 supports tumour progression in vivo despite having limited effects on tumour cell proliferation in vtiro. To determine whether these effects require host immune competence, we inoculated *Pcif1*-deficient and shNTC control Yumm1.7, BP and LLC1 cells into athymic *Balb/cNj-Foxn1^nu^/Gpt* mice, which lack functional T and B cells owing to defective thymic epithelium development (**Extended Data Fig 1c**). Under these conditions, PCIF1 loss no longer affected tumour growth in any models, indicating that the anti-tumour effects elicited by tumour-intrinsic PCIF1 depletion requires a functional adaptive immune system (**Extended Data Fig 1c**).

**Figure 1.**
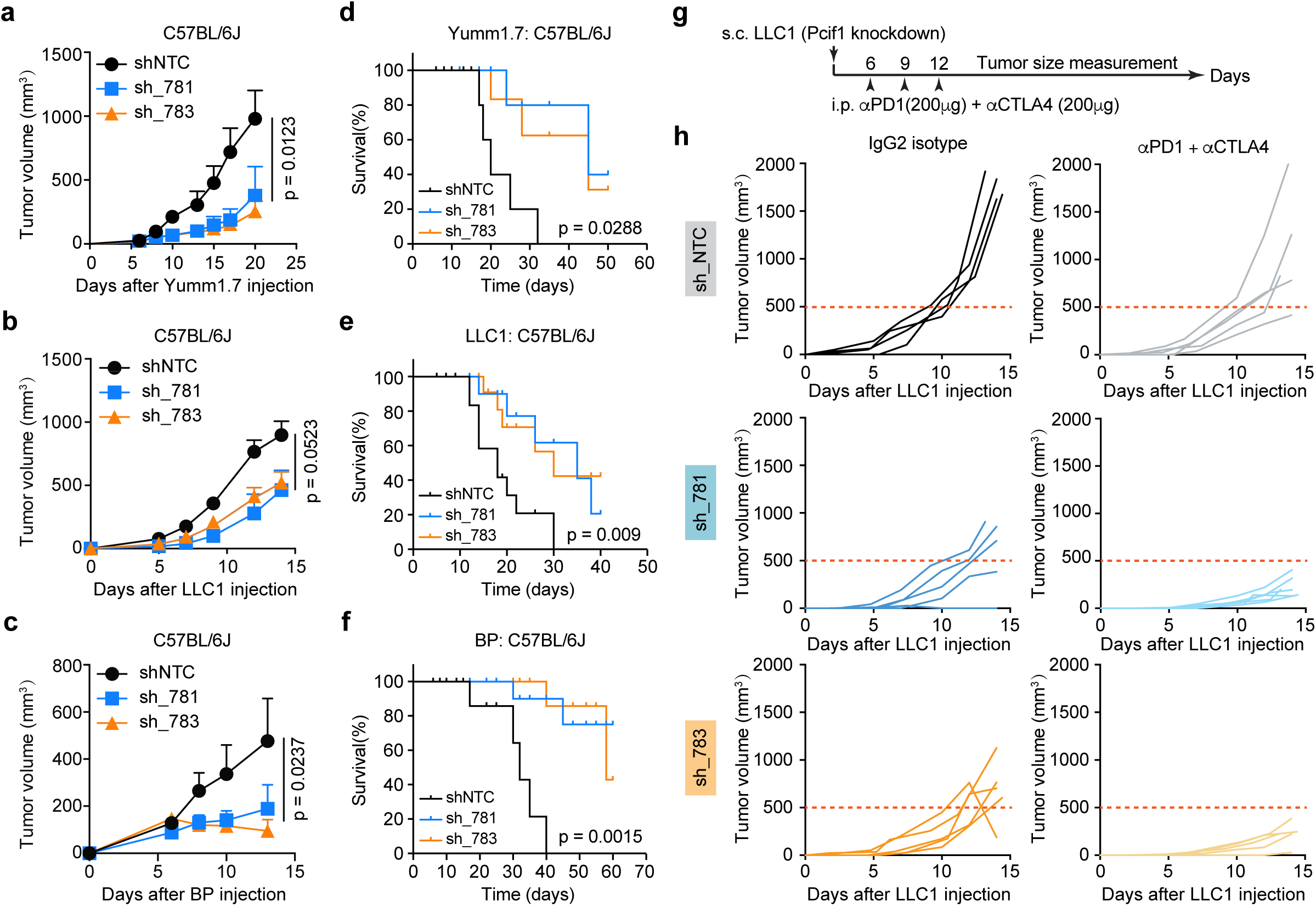
Tumour-cell PCIF1 loss restrains syngeneic tumour growth and sensitizes LLC1 tumours to checkpoint blockade. (**a-f**) In vivo growth and survival of syngeneic tumours generated from control or Pcif1-depleted tumour cells in immunocompetent C57BL/6J mice. Yumm1.7 melanoma, LLC1 lung carcinoma and BP melanoma cells were transduced with a non-targeting shRNA (shNTC) or two independent Pcif1 shRNAs (sh_781 and sh_783) before subcutaneous implantation. Mean tumour-growth curves are shown for Yumm1.7 (**a**; n = 5 mice per group), LLC1 (**c**; n = 5-8 mice per group) and BP (**e**; n = 5 mice per group) tumours. Kaplan–Meier survival curves for the corresponding cohorts are shown in **b**, **d** and **f**, respectively. (**g**) Experimental design for immune checkpoint blockade in the LLC1 model. Mice were implanted subcutaneously with shNTC-, sh_781- or sh_783-expressing LLC1 cells and treated intraperitoneally with anti-PD1 antibody (200 μg) plus anti-CTLA4 antibody (200 μg) on days 6, 9 and 12 after tumour implantation. Tumour size was measured longitudinally. **(h)** Individual tumour-growth curves of LLC1 tumours expressing shNTC, sh_781 or sh_783 in mice treated with IgG2 isotype control or combined anti-PD1 plus anti-CTLA4 therapy; n = 4-6 mice per group. Red dashed lines indicate a tumour volume of 500 mm³. Data in **a**, **c** and **e** are shown as mean ± s.e.m. P values for tumour growth curves were calculated by two-way ANOVA. P values for survival curves were calculated by log-rank test.

### PCIF1 knockdown sensitizes tumours to ICB

To evaluate whether perturbation of PCIF1 could synergize with ICB-based therapy^19^, we conducted *in vivo* studies using syngeneic mouse models inoculated with *Pcif1*-deficient LLC1 lung carcinoma or Yumm1.7 melanoma cells. Mice were treated with anti-PD1 antibody alone or in combination with anti-CTLA4 antibody (**Fig 1g and Extended Data Fig 2a**). In the LLC1 model, combined anti-PD1 and anti-CTLA4 treatment had minimal and inconsistent effects on tumour growth (**Fig 1h**), while anti-PD1 monotherapy failed to elicit significant tumour control in the Yumm1.7 model (**Extended Data Fig 2b**). These results are consistent with the well-documented immune-refractory phenotypes of both tumour models. LLC1 cells, derived from a spontaneous Lewis lung carcinoma in C57BL/6J mice and serially passaged in immunocompetent hosts^18^, are known to be highly immune evasive and resistant refractory to ICB therapy^20^; similarly, Yumm1.7 cells have previously been shown to proliferate despite ICB treatment^21^. However, Pcif1 knockdown alone significantly reduced tumour growth in LLC1-bearing mice, even in the absence of ICB (**Fig 1h**), in line with our previous observations. Moreover, the combination of Pcif1 knockdown with treatment of anti-PD1 plus anti-CTLA4 resulted in a marked and synergistic suppression of LLC1 tumour growth (**Fig 1h**). A comparable synergistic effect was also observed in the otherwise ICB-refractory Yumm1.7 model (**Extended Data Fig 2b**), underscoring the potential of PCIF1 inhibition to overcome tumour intrinsic resistance to immunotherapy.

### PCIF1 depletion induces tumour inflammation

The immune dependence of tumour control prompted us to ask whether tumour cell PCIF1 loss alters the inflammatory state of the tumour microenvironment. Immunohistochemical analysis confirmed sustained depletion of PCIF1 protein in LLC1 tumours (**Extended Data Fig. 3a**). Bulk RNA sequencing of these tumours derived from shNTC and sh_*Pcif1* groups revealed a pronounced enrichment of immune- and inflammation-associated pathways upon *Pcif1* depletion, as assessed by single-sample gene set enrichment analysis (ssGSEA) (**Extended Data Fig. 3b**). Among the most consistently enriched pathways were inflammatory response (**Extended Data Fig. 3c**, top), interferon-γ response (**Extended Data Fig. 3c**, bottom) and TNFα signalling (**Fig. 2a**), indicating that tumour cell PCIF1 loss is accompanied by a shift toward an inflamed tumour state *in vivo*.

**Figure 2.**
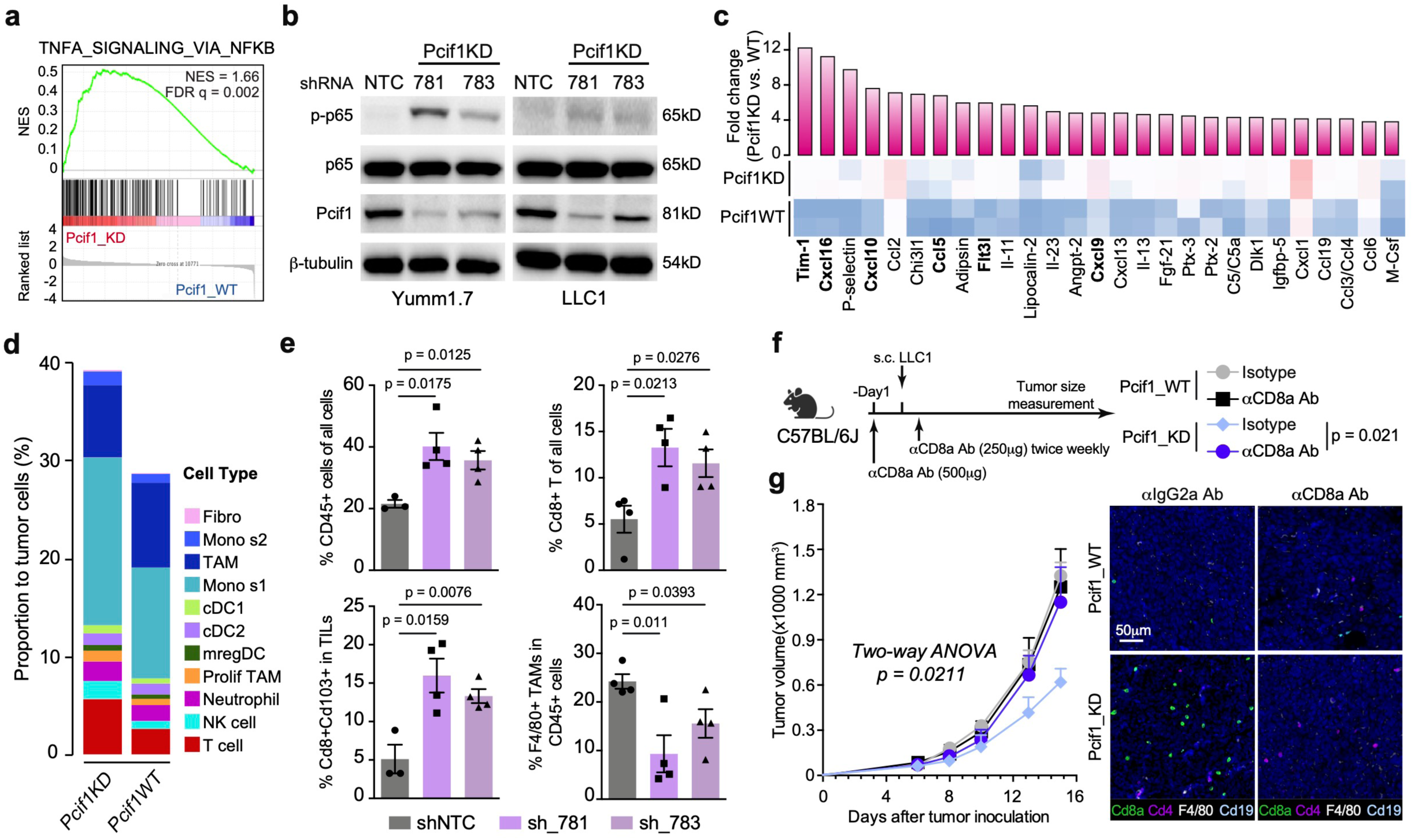
PCIF1 loss induces tumour inflammation and CD8□ T-cell-dependent tumour control. **(a)** Gene set enrichment analysis of bulk RNA-seq data from control and Pcif1-depleted LLC1 tumours showing enrichment of the TNFα signalling via NFκB pathway in Pcif1-depleted tumours. NES and FDR q-value are indicated. **(b)** Immunoblot analysis of phosphorylated p65, total p65 and PCIF1 in Yumm1.7 and LLC1 tumour cells expressing a non-targeting shRNA (NTC) or two independent Pcif1 shRNAs (sh_781 and sh_783). β-tubulin was used as a loading control. Blots are representative of n = 3 independent experiments with similar results. **(c)** Cytokine proteome array analysis of conditioned medium from control and Pcif1-depleted LLC1 cells. Bars show fold change in cytokine abundance in Pcif1-depleted cells relative to control cells; the heatmap indicates relative abundance in the corresponding conditions. **(d)** Single-cell RNA-seq-based cell-type composition of LLC1 tumours generated from control or Pcif1-depleted cells. Bars show the relative abundance of annotated immune and stromal populations normalized to tumour cells. **(e)** Flow-cytometric quantification of tumour-infiltrating immune populations in LLC1 tumours expressing shNTC, sh_781 or sh_783. Plots show the percentage of CD45□ cells among all cells, CD8□ T cells among total cells, CD8□CD103□ cells among tumour-infiltrating lymphocytes and F4/80□ tumour-associated macrophages among CD45□ cells; n = 4 biologically independent tumours per group. **(f)** Experimental scheme for CD8□ T-cell depletion in the LLC1 model. C57BL/6J mice were implanted subcutaneously with control or Pcif1-depleted LLC1 cells and treated with anti-CD8a antibody or isotype control according to the indicated schedule. Tumour size was monitored over time. **(g)** Tumour-growth curves of control and Pcif1-depleted LLC1 tumours in mice treated with anti-CD8a antibody or isotype control, together with representative multiplex immunofluorescence images showing CD8a, CD4, F4/80 and CD19 staining in tumour sections; n = 5 mice per group. Scale bar, 50 μm. Data in **e** and **g** are shown as mean ± s.e.m. P values in **e** were calculated by two-sided unpaired t-test. P values for tumour-growth curves in **g** were calculated by two-way ANOVA.

To determine whether this transcriptional inflammatory state was reflected at the signalling level, we examined TNFα signalling at the protein level. PCIF1 depletion increased phosphorylation of p65 in both LLC1 and Yumm1.7 cells (**Fig. 2b**). Histopathological analysis of *Pcif1*-deficient tumours further revealed increased vascular remodelling by haematoxylin and eosin staining (**Extended Data Fig. 3a**, yellow arrowheads), in agreement with the enrichment of angiogenesis-related gene sets observed by ssGSEA (**Extended Data Fig. 3b**). As tumour-draining lymph nodes (tdLNs) represent a key site for the priming of adaptive anti-tumour immune responses, we next profiled ipsilateral tdLNs from tumour-bearing mice by RNA sequencing (**Extended Data Fig. 3a and 3d**). Pathway analysis revealed coordinated enrichment of immune-related signatures (**Extended Data Fig. 3d**), including interferon-α responsive signalling, inflammatory response, interferon-γ responsive signalling and NFκB signalling (**Extended Data Fig 3e**). These data indicate that tumour cell PCIF1 depletion is associated not only with local tumour inflammation, but also with immune activation in the draining lymphoid compartment.

**Figure 3.**
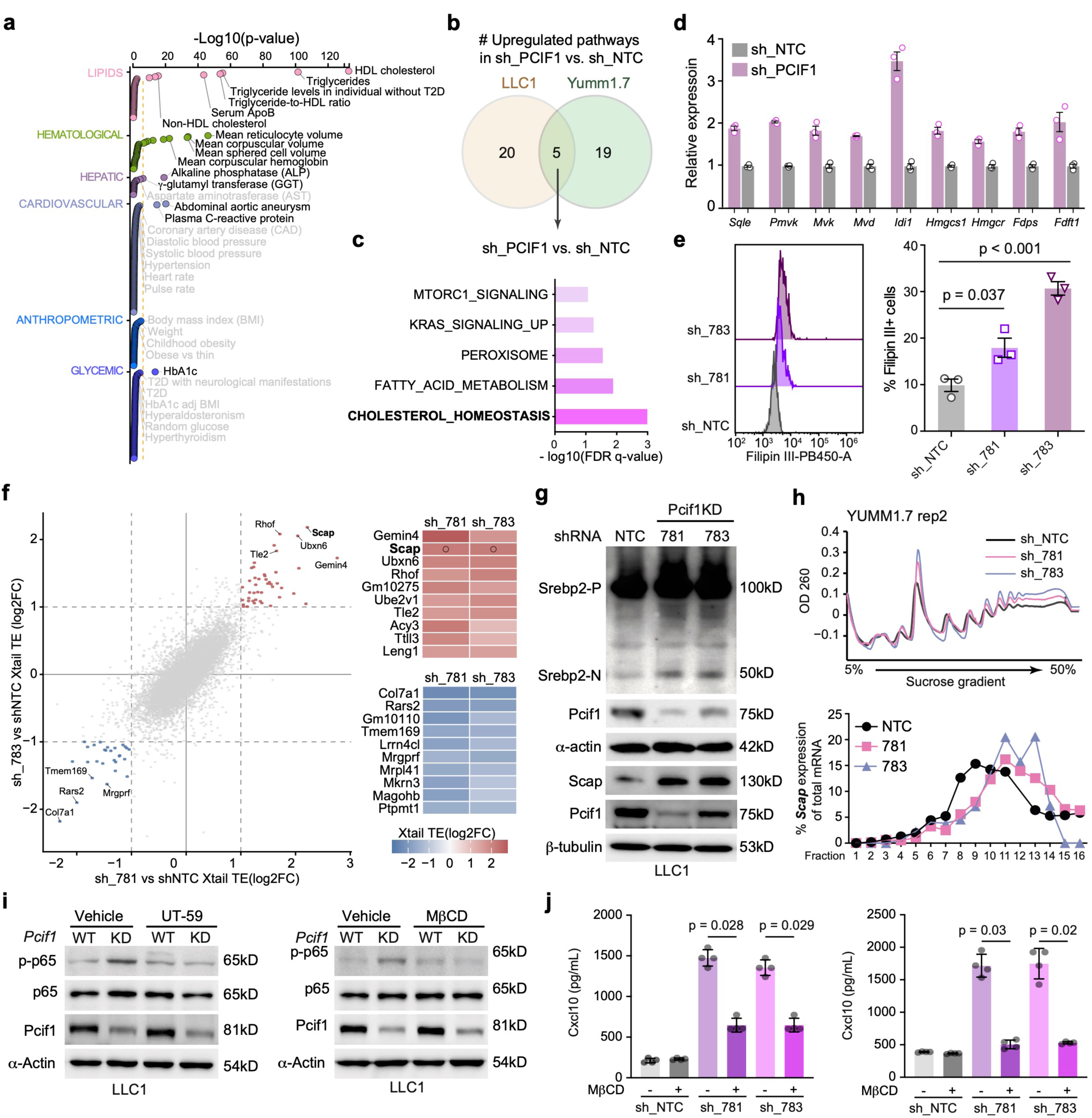
PCIF1 restrains SCAP-dependent cholesterol-linked inflammatory signalling. **(a)** Human genetic evidence linking PCIF1 to metabolic traits. Associations between PCIF1 loss-of-function genetic variation and human disease or metabolic phenotypes were analysed using the Common Metabolic Diseases Knowledge Portal. Traits are grouped by biological category, and association strength is shown as −log10(P value). **(b)** Overlap of pathways upregulated in Pcif1-depleted LLC1 and Yumm1.7 tumour cells relative to non-targeting controls, as determined by RNA-seq pathway enrichment analysis. **(c)** Shared upregulated pathways in Pcif1-depleted LLC1 and Yumm1.7 cells ranked by enrichment significance, highlighting cholesterol homeostasis as the top shared programme. **(d)** Quantitative PCR analysis of cholesterol-biosynthetic genes in control and Pcif1-depleted tumour cells; n = 3 biologically independent samples per group. **(e)** Flow-cytometric analysis of free cholesterol using filipin III staining in LLC1 cells expressing shNTC, sh_781 or sh_783; n = 3 biologically independent samples per group. Representative histograms are shown on the left, and quantification of filipin III-positive cells is shown on the right. **(f)** Xtail analysis of polysome RNA-seq data from LLC1 cells after Pcif1 depletion. Total input RNA and polysome-associated RNA were profiled by RNA-seq in LLC1 cells expressing a non-targeting shRNA (shNTC) or two independent Pcif1 shRNAs (sh_781 and sh_783), and changes in translational efficiency were inferred using xTAIL. The scatter plot shows transcript-level translational-efficiency changes in sh_781 versus shNTC cells on the x axis and sh_783 versus shNTC cells on the y axis. Each dot represents one transcript. Red dots indicate transcripts with increased translational efficiency in both Pcif1 knockdown conditions, and blue dots indicate transcripts with decreased translational efficiency in both Pcif1 knockdown conditions, using the indicated log□ fold-change thresholds. Selected recurrently regulated transcripts are labelled. The heat maps show the top shared transcripts with increased translational efficiency (top) or decreased translational efficiency (bottom) in both Pcif1 knockdown conditions. Scap was identified among the shared transcripts with increased translational efficiency after Pcif1 depletion. **(g)** Immunoblot analysis of SREBP2 precursor (SREBP2-P), cleaved N-terminal SREBP2 (SREBP2-N), PCIF1 and SCAP in control and Pcif1-depleted LLC1 cells. β-tubulin and α-actin were used as loading controls. Blots are representative of n = 2 independent experiments with similar results. **(h)** Polysome profiling of Yumm1.7 cells expressing shNTC, sh_781 or sh_783 on a 5–50% sucrose gradient, with qPCR quantification of Scap mRNA distribution across gradient fractions. Data are representative of n = 3 independent experiments with similar results. **(i)** Immunoblot analysis of phosphorylated p65, total p65 and PCIF1 in LLC1 control and Pcif1-depleted cells treated with vehicle, the SCAP inhibitor UT-59 or the cholesterol-depleting agent methyl-β-cyclodextrin (MβCD). α-actin was used as a loading control. Both immunoblot panels in h were performed in LLC1 cells. Blots are representative of n = 2 independent experiments with similar results. **(j)** ELISA quantification of CXCL10 secretion from LLC1 cells (top) and Yumm1.7 cells (bottom) expressing shNTC, sh_781 or sh_783 and treated with vehicle or MβCD; n = 4 biologically independent samples per group. Data in d, e and i are shown as mean ± s.e.m. P values in e and i were calculated by two-sided unpaired t-test.

We then asked whether the inflammatory transcriptional state induced by PCIF1 loss was accompanied by altered secretion of immune-modulatory factors from tumour cells. Secretome profiling of LLC1 tumour cells using a mouse XL cytokine proteome array (**Extended Data Fig. 3f**) identified increased abundance of several inflammatory and immune-recruting factors after Pcif1 depletion, including T cell-recruiting chemokines Cxcl9 and Cxcl10^22^, the NK- and T cell-associated factors Tim-1^23^ and Cxcl16^24^, and the dendritic cell differentiation factor Flt3l^25^ (**Fig. 2c**). Cxcl10 induction was further validated by ELISA in two independent tumour cell models (**Extended Data Fig 3g**), supporting the idea that PCIF1 loss promotes a tumour cell secretory programme capable of recruiting and shaping immune infiltrates. Motivated by these observations, we next assessed immune cell composition at the tumour and tdLN level by deconvoluting bulk RNA-sequencing data using CIBERSORTx^26^. Pcif1-depleted tumours showed increased inferred fractions of CD8+ T cells and dendritic cell populations, whereas tdLNs showed increased T cell and dendritic cell signatures (**Extended Data Fig. 3h**), consistent with enhanced immune activation within tumour and lymphoid tissue.

### CD8+ T cells mediate PCIF1-loss tumour control

To define the immune changes induced by tumour cell PCIF1 at single-cell resolution, we performed unbiased single-cell RNA sequencing (scRNA-seq) to profile immune infiltration in LLC1 tumours. Graph-based clustering identified the major cellular compartments within the tumour, with malignant epithelial cells constituting the predominant population, as well as fibroblasts, T cells, NK cells, neutrophils and multiple myeloid populations, including monocytes, tumour-associated macrophages (TAMs) and dendritic-cell substes (**Extended Data Fig 4a-b**). Malignant epithelial cells lacked expression of the pan-leukocyte marker *Ptprc* and expressed tumour-associated genes including *Kras*, *Cav1, Mapk1*, *Igf2bp3*, *Agpat3* and *Agpat5* (**Extended Data Fig 4c-d**). Their malignant identity was further supported by inferCNV analysis^27^, which revealed broad chromosome-scale copy number alterations in the annotated cancer cell compartment, in contrast to immune and stromal cells with largely neutral copy number profiles (**Extended Data Fig. 4e**). The remaining immune and stromal compartments were annotated using canonical lineage markers and established single cell immune-state definitions, including monocyte, neutrophil, TAM, cDC1, cDC2 and mregDC populations^28^ (**Extended Data Fig. 4c-d**). Since we have observed the broad inflammatory remodelling and NFκB activation upon Pcif1 loss (**Extended Data Fig 3**), we examined the GAS/STING and innate immune pathway expression by western blot. However, we did not observe any significant alterations of related proteins, including TBK1, STAT1, IRF3 and STING upon Pcif1 knockdown (**Extended Data Fig. 4f**).

Consistent with the reduced tumour outgrowth observed *in vivo*, malignant epithelial cells expressing the proliferation marker *Mki67* were more abundant in shNTC tumours than in tumours expressing either of two independent Pcif1 shRNAs (**Extended Data Fig. 5a**). By contrast, most myeloid and stromal compartments showed relatively modest changes in abundance. The most prominent immune alteration was an increase in T cells in Pcif1-depleted tumours, accompanied by a modest reduction in TAMs (**Fig. 2d and Extended Data Fig. 4b**). This T cell-enriched state was independently validated by multiplexed immunofluorescence (mIF) in LLC1 tumours, which showed increased infiltration of Cd8a^+^ T cells in *Pcif1*-deficient tumours (**Extended Data Fig. 5b**). Flow cytometric analysis further confirmed increased CD45^+^ immune infiltration, largely attributable to CD8^+^ T cells, in tumours lacking Pcif1 (**Fig. 2e** and **Extended Data Fig. 5c**). We next examined whether PCIF1 loss altered the functional composition of tumour infiltrating T cells. Subclustering of T cells resolved CD4^+^ T cells, γδ T cells and multiple CD8+ T cell states, including cytotoxic CD8^+^ T cells, progenitor-like exhausted T cells (CD8Tpex), terminally exhausted T cells (CD8Tex), interferon-stimulated gene (ISG)^+^ T cells (ISG+ CD8T), effector memory T cells (CD8Tem) and naïve-like CD8^+^ T cells^29^ (**Extended Data Fig. 6a**, left **and 6b**). Pcif1-depleted tumours showed an increased fraction of cytotoxic CD8+ T cells, whereas exhausted CD8+ T cells and ISG+ CD8 T cell states were significantly reduced (**Extended Data Fig. 6c**). At functional marker level, T cells from Pcif1-depleted tumours expressed higher levels of the progenitor-associated transcription factor *Tcf7* and the activation marker *Cd69*, alongside lower levels of exhaustion-associated genes including *Tox*, *Pdcd1*, *Entpd1* and *Havcr2* (**Extended Data Fig. 6d**). CD4^+^ T cell analysis further suggested reduced regulatory T cells (Treg) features (**Extended Data Fig. 6c**), including the decreased *Foxp3* expression and lower expression of *Ctla4* and *Lag3* (**Extended Data Fig 6e**). Consistent with these transcriptional changes, flow cytometry showed an increased frequency of activated effector Cd8a^+^Cd103^+^ T cells and a reduction in F4/80^+^ TAMs following PCIF1 knockdown (**Fig 2e**).

To functionally assess the contribution of CD8^+^ T cells to tumour control, we depleted CD8^+^ T cells with an anti-Cd8a antibody beginning at tumour implantation (**Fig. 2f**). Antibody treatment efficiently reduced intra-tumoural CD8^+^ T cells, as confirmed by mIF, without affecting CD4^+^ T cell abundance (**Fig. 2g**, right). Notably, CD8^+^ T cell depletion largely restored the growth of Pcif1-deficient LLC1 tumours to levels comparable to control tumours (**Fig. 2g**, left). Thus, the tumour growth restraint induced by tumour cell PCIF1 loss is mediated, at least in major part, by CD8^+^ T cell immunity.

### PCIF1 loss induces cholesterol biosynthesis

Having established that tumour cell PCIF1 loss induces a CD8+ T cell-dependent antitumour response, we next sought to identify the tumour intrinsic programme that links PCIF1 to inflammatory immune activation. PCIF1 is a cap-specific mRNA methyltransferase that deposits m^6^Am at the 5′ cap^2^. In viral infection models, PCIF1-dependent cap methylation can promote immune evasion by regulating viral transcript stability and translation, thus attenuating interferon responses^13,15^, In cancer immunity, PCIF1 has recently been implicated in CD8^+^ T cell intrinsic regulation of effector activity and ferroptosis^30^. These observations raised the possibility that PCIF1 might also shape antitumour immunity through a tumour cell intrinsic post-transcriptional mechanism.

Human genetic data provided an unexpected clue. Genome-wide association studies (GWAS) showed that loss-of-function variants in PCIF1 were associated with lipid and metabolic traits, including plasma cholesterol, triglycerides, and the triglyceride-to-high-density lipoprotein (HDL) ratio (**Fig. 3a**). These associations were consistent with previously reported links between PCIF1 and increased body mass index (BMI) and diabete-related traits^31^ (**Extended Data Fig. 7a**), suggesting that PCIF1 may participate in lipid metabolic regulation. We therefore asked whether PCIF1 depletion altered translation or metabolic gene expression in tumour cells. Polysome profiling in LLC1 and Yumm1.7 cells revealed no major change in global translation after Pcif1 knockdown (**Extended Data Fig. 7b**). Polysome-associated RNA-seq further showed that genome-wide translational efficiency remained largely unchanged (**Extended Data Fig. 7c**), indicating that PCIF1 loss does not broadly modulate general mRNA translation in these models^32^. We thus examined transcriptional changes induced by PCIF1 depletion. The RNA-seq of LLC1 and Yumm1.7 cells identified a small set of pathways consistently upregulated upon Pcif1 knockdown, with cholesterol homeostasis emerging as the most enriched shared programme (**Fig. 3b-c** and **Extended Data Fig. 7d**). Quantitative PCR confirmed increased expression of cholesterol biosynthetic genes, including *Hmgcr*, *Mvk*, and *Sqle* (**Fig. 3d and Extended Data Fig. 7e**), suggesting coordinated activation of the cholesterol biosynthesis pathway. We then tested whether this transcriptional programme is translated into altered cellular cholesterol content. Filipin III staining, a fluorescent polyene that binds free cholesterol^33^, revealed increased cholesterol signal in LLC1 cells expressing independent Pcif1 shRNAs evaluated by flow cytometry (**Fig. 3e and Extended Data Fig. 7f**). Confocal imaging further showed increased cytoplasmic filipin III-positive structures that colocalized with lipid droplets labelled with BODIPY C16, consistent with increased intracellular cholesterol and altered lipid storage (**Extended Data Fig. 7g**). Thus, PCIF1 depletion activates a cholesterol biosynthetic state that is reflected at both transcriptional and cellular lipid levels.

To obtain a broader view of lipid remodelling, we performed untargeted lipidomic profiling using liquid chromatography-mass spectrometry (LC-MS/MS) (**Extended Data Fig. 8a**). Representative total ion chromtograms (TICs) in both ionization modes confirmed stable LC-MS signal acquisition and chromatographic separation across quality control sample (QC) and biological samples, supporting the overall quality of the metabolomics dataset (**Extended Data Fig. 8b**). After intensity normalization (**Extended Data Fig. 8c-d**), partial least squares discriminant analysis (PLS-DA) clearly separated *Pcif1*-proficient and *Pcif1*-deficient cells (**Extended Data Fig. 8e**). Consistent with the transcriptional enrichment of cholesterol and lipid metabolic programs, LC-MS lipidomics revealed a broad reorganization of the cellular lipidome following PCIF1 depletion. Differential abundance analysis in both negative and positive ion modes identified coordinated changes in lipid species that clearly distinguished *Pcif1*-deficient from *Pcif1*-proficient cells (**Extended Data Fig. 8f-g**). At the lipid class level, PCIF1 loss was associated with an increase in PE species and a marked reduction in PG and cardiolipin-related lipids in negative mode, while positive-mode profiling highlighted changes in PC, SM, GlcCer and TAG species (**Extended Data Fig. 8h-i**). These coordinated transcriptional and metabolic changes indicate that PCIF1 depletion reshapes tumour lipid metabolism, promoting intracellular cholesterol biosynthesis and broader membrane lipid remodelling.

### SCAP translation promotes cholesterol-linked inflammation

Although PCIF1 depletion did not broadly alter global translation, we next asked whether it selectively modulated the translational engagement of specific transcripts that may impact on the transcription of cholesterol-related pathway genes. We analysed polysome-associated RNA-seq data using Xtail^34^. In LLC1 cells, 44 transcripts showed increased translational efficiency and 28 transcripts showed decreased translational efficiency in both independent Pcif1 knockdown conditions (**Fig. 3f**, left and **Extended Data Fig. 9a-b**). Among these candidates, we focused on sterol regulatory element-binding protein cleavage-activating protein (SCAP), a central regulator of cholesterol sensing and SREBP2 activation^35^, because Scap showed increased translational engagement after PCIF1 depletion (**Fig. 3f**, right). A similar subset of translationally regulated transcripts was identified in Yumm1.7 cells, particularly Scap transcript (**Extended Data Fig. 9a-b**).

Under cholesterol-depleted conditions, SCAP escorts SREBP2 from the endoplasmic reticulum to the Golgi apparatus, where SREBP2 is proteolytically processed to generate the transcriptionally active N-terminal fragment, SREBP2-N, that induces cholesterol biosynthetic genes^36^ (**Extended Data Fig. 9c**, left). This regulatory architecture provided a plausible route by which selective translational control could amplify cholesterol biosynthesis after PCIF1 loss. In accordance, PCIF1 depletion increased SCAP protein abundance and elvated the cleaved SREBP2-N fragment in both LLC1 and Yumm1.7 tumour cells (**Fig. 3g and Extended Data Fig. 9d**). By contrast, the mRNA levels of *Scap* and other components of the SCAP-SREBP2 trafficking machinery, including *Sec24a*, *Insig2*, *S1p*, *S2p* and *Sar1a,* were not substantially altered (**Extended Data Fig. 9c, right**). Polysome fraction qPCR further confirmed a redistribution of *Scap* transcripts toward heavier polysome fractions in PCIF1 depleted Yumm1.7 (**Fig 3h**), whereas the control transcript *Hprt1* showed no comparable shift (**Extended Data Fig. 9e**). Similar enrichment of polysome-associated *Scap* transcripts was observed in LLC1 (**Extended Data Fig. 9f-h**). Thus, PCIF1 loss selectively enhances SCAP translational engagement, providing a post-transcriptional regulatory point for SREBP2-dependent cholesterol biosynthesis.

We next asked whether this SCAP-cholesterol axis contributes to the inflammatory signalling induced by PCIF1 loss. Pharmacological inhibition of SCAP with UT-59, which interferes with SCAP-dependent SREBP2 activation^37^, effectively suppressed the increase in p65 phosphorylation observed in PCIF1-depleted tumour cells (**Fig. 3i**, left and **Extended Data Fig. 9i**). Similary, acute cholesterol depletion with methyl-β-cyclodextrin (MβCD)^38,39^ reduced p65 phosphorylation in both cell models (**Fig 3i**, right and **Extended Data Fig. 9j**). Conversely, treatment with U18666A, an inhibitor of NPC1-dependent lysosomal cholesterol export that promotes intracellular cholesterol accumulation^40^, further increased phosphorylation of p65 in PCIF1-deficient cells (**Extended Data Fig. 9k**). These pharmacological perturbations suggest that intracellular cholesterol contribute to TNFα pathway activation after PCIF1 loss. Furthermore, we tested whether cholesterol availability was required for the chemokine output of this inflammatory state. MβCD treatment reduced the elevated secretion of Cxcl10 from PCIF1-deficient tumour cells (**Fig. 3j and Extended Data Fig. 9l**), linking cholesterol dependent signalling to a T cell recruiting chemokine induced by PCIF1 loss. Together, these data support a model in which PCIF1 restrains SCAP translation engagement, when this retraint is lost, SCAP-SREBP2-dependent cholesterol biosynthesis promotes an inflammatory programme in tumour cells.

### PCIF1 marks poor immune checkpoint response

Given that tumour-cell PCIF1 loss enhanced CD8□ T-cell-dependent tumour control and sensitized syngeneic tumours to checkpoint blockade, we next asked whether PCIF1 expression was associated with clinical outcome in patients treated with ICB. Although established biomarkers, such as tumour mutational burden (TMB)^41^, PD-L1 expression^42^ and intra-tumoral CD8□ T-cell infiltration^43^, have been shown to correlate with ICB efficacy, there remains a substantial unmet need to identify additional, potentially modifiable tumour-intrinsic factors that may influence therapeutic outcomes.

We first analysed melanoma cohorts treated with anti-PD1 therapy. In this integrated cohort, high PCIF1 expression was associated with shorter overall survival and progression-free survival when patients were stratified using the available tumour transcriptomes (**Fig. 4a** and **Extended Data Fig. 10a**). We then examined whether this association was already present before therapy or became more apparent during treatment. In pre-treatment biopsies, high PCIF1 expression was associated with shorter overall survival compared with low PCIF1 expression (HR = 1.48, 95% CI 1.04–2.10; log-rank P = 0.027; **Fig. 4a,b**). Consistent with this survival difference, PCIF1 expression was higher in pre-treatment samples from non-responders than from responders, as defined by RECIST 1.1 criteria (**Fig. 4b**, right). In on-treatment biopsies from the same anti-PD1 melanoma cohort, stratification by PCIF1 expression produced a stronger separation of survival curves, with high PCIF1 expression associated with inferior overall survival (HR = 2.24, 95% CI 1.12–4.50; log-rank P = 0.019; **Fig. 4a,c**). Non-responders also showed higher PCIF1 expression in on-treatment samples (**Fig. 4c**, right), indicating that sustained or treatment-associated PCIF1 expression is linked to reduced clinical benefit.

**Figure 4.**
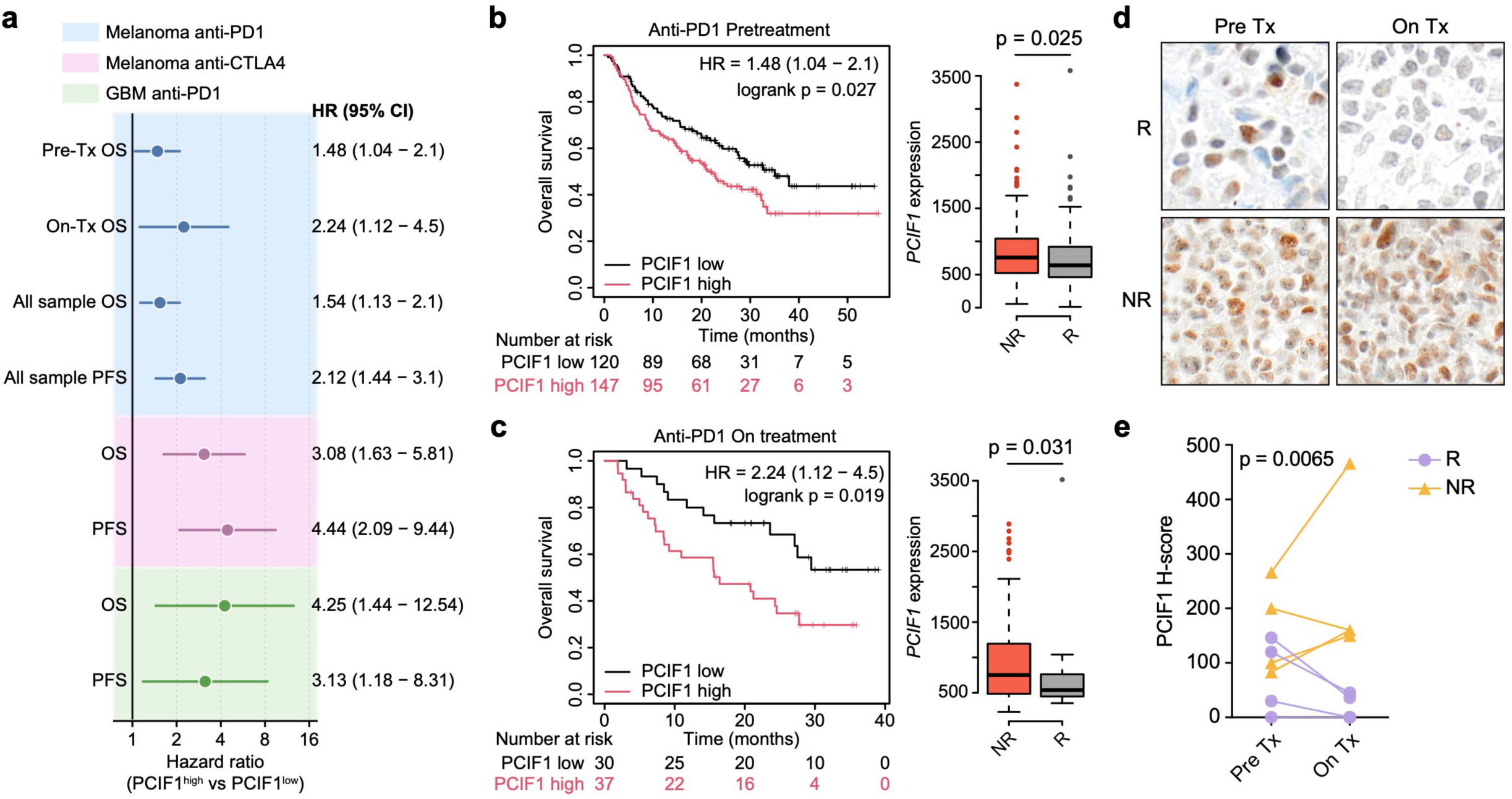
High PCIF1 expression marks reduced benefit from immune checkpoint blockade. **(a)** Forest plot showing the association between PCIF1 expression and clinical outcome in ICB-treated patient cohorts. Hazard ratios (HRs) and 95% confidence intervals are shown for PCIF1-high versus PCIF1-low groups defined using the optimal expression cutoff selected by KM Plotter^44^ and plotted on a logarithmic scale. HR values greater than 1 indicate worse outcome in the PCIF1-high group. The melanoma anti-PD1 analyses include complementary stratifications of the same clinical dataset, including pre-treatment overall survival (Pre-Tx OS), on-treatment overall survival (On-Tx OS), and overall OS and progression-free survival (PFS) analyses using the available anti-PD1-treated samples. Additional treatment contexts include melanoma patients treated with anti-CTLA4 and glioblastoma patients treated with anti-PD1. **(b)** Kaplan-Meier analysis of overall survival in anti-PD1-treated melanoma patients stratified by PCIF1 expression in pre-treatment tumour biopsies; The corresponding box plot shows PCIF1 expression in pre-treatment biopsies from responders (R) and non-responders (NR); n = 183 R and n = 232 NR samples. **(c)** Kaplan-Meier analysis of overall survival in anti-PD1-treated melanoma patients stratified by PCIF1 expression in on-treatment tumour biopsies; The corresponding box plot shows PCIF1 expression in on-treatment biopsies from responders and non-responders; n = 183 R and n = 232 NR samples. **(d)** Representative immunohistochemical staining of PCIF1 in paired pre-treatment and on-treatment tumour biopsies from melanoma patients receiving nivolumab, shown for representative responders and non-responders. Scale bar, 20 μm. **(e)** Quantification of PCIF1 immunohistochemical staining by H-score in paired pre-treatment and on-treatment melanoma biopsies from responders and non-responders; Each line represents one patient. In **b** and **c**, survival differences were assessed by log-rank test. P values for response-group comparisons in the box plots in **b** and **c** were calculated by Mann–Whitney test. P value in **e** was calculated by two-sided t-test.

We next asked whether the association between high PCIF1 expression and poor ICB outcome extended to other treatment settings and tumour types. In melanoma patients treated with anti-CTLA4 therapy, high PCIF1 expression was associated with inferior overall survival and progression-free survival (**Fig. 4a and Extended Data Fig. 10b**). A similar association was observed in glioblastoma patients receiving anti-PD1 therapy, in whom high PCIF1 expression was linked to shorter overall survival and progression-free survival (**Fig. 4a and Extended Data Fig. 10c,d**). Thus, although the anti-PD1 melanoma analyses represent complementary stratifications of the same clinical dataset, the overall pattern was also observed in distinct ICB contexts, including anti-CTLA4-treated melanoma and anti-PD1-treated glioblastoma.

To assess PCIF1 expression at the tissue level, we performed immunohistochemical analysis of paired pre-treatment and on-treatment tumour biopsies from melanoma patients receiving nivolumab. Non-responders generally exhibited higher PCIF1 staining at baseline and maintained or increased PCIF1 expression during treatment (**Fig. 4d,e**). By contrast, responders tended to show lower baseline PCIF1 expression and stable or reduced PCIF1 staining after treatment (**Fig. 4d,e**). Although limited by sample size, this in-house tissue analysis was concordant with the transcriptomic association between elevated PCIF1 expression and reduced ICB benefit. Finally, we examined whether high PCIF1 expression was associated with reduced cytotoxic immune activity in patient tumours. Across multiple tumour types, including glioblastoma, lung adenocarcinoma, breast cancer and advanced melanoma, PCIF1 expression inversely correlated with cytolytic T-lymphocyte scores (**Extended Data Fig. 10e**). Additionally, high PCIF1 expression was also associated with inferior overall survival across several cancer types, including lung adenocarcinoma, colon cancer, ER-positive breast cancer and serous ovarian cancer (**Extended Data Fig. 10f,g**). These latter analyses are not ICB-specific, but they support a broader association between elevated PCIF1 expression, reduced cytotoxic immune activity and adverse clinical outcome. Together with the preclinical data, these findings indicate that PCIF1 marks reduced ICB therapy benefit and is associated with a less cytotoxic tumour immune context in patients.

## Discussion

This study identifies tumour cell PCIF1 as a checkpoint on the immunological consequences of cholesterol biosynthesis. PCIF1 loss uncovered an inflammatory state *in vivo*, built around SCAP translational engagement, SREBP2-dependent cholesterol biosynthesis, NFκB activation and production of T cell-recruiting chemokines. This state was accompanied by CD8□ T cell infiltration, CD8□ T cell-dependent tumour control and improved response to ICB. Together, these findings support a model in which PCIF1 restrains a cholesterol-coupled inflammation axis that can render tumours more visible to adaptive immunity when this restraint is removed (**Extended Data Fig. 10h**). These data do not argue that cholesterol biosynthesis is intrinsically favourable or unfavourable for antitumour immunity. Rather, they suggest that a metabolic pathway usually associated with membrane production, metabolic fitness and tumour growth can, in the right regulatory context, acquire an inflammatory output. The relevant question may therefore be less whether cholesterol metabolism is pro- or antitumoural, but more how tumour cells transmit cholesterol-linked signals to their microenvironment. PCIF1 may provide one such gate, by limiting SCAP-dependent cholesterol biosynthesis and its downstream inflammatory output, PCIF1 helps keep a growth-associated metabolic programme from becoming a CD8□ T cell-recruiting tumour state.

This model places cap-proximal RNA regulation upstream of metabolic immune control. PCIF1 has been linked to viral immune evasion and to T cell-intrinsic regulation of antitumour immunity, but our data position PCIF1 within tumour cells as a regulator of lipid metabolism and immune visibility. The present evidence supports selective SCAP translational engagement after PCIF1 loss, rather than a global remodelling of translation.

Whether SCAP is a direct m□Am-regulated PCIF1 substrate remains to be resolved by transcript-specific m□Am mapping and catalytic rescue. Similarly, the connection between cholesterol accumulation and NFκB activation is likely to involve additional lipid-sensing or stress-response intermediates. These open questions define the next mechanistic layer of this axis. From clinical viewpoint, high PCIF1 expression was associated with reduced benefit in ICB-treated melanoma and glioblastoma cohorts and with lower cytolytic T-cell signatures across tumour types. These retrospective associations are concordant with the experimental finding that PCIF1 restrains CD8□ T cell-mediated tumour control. Therapeutically, our work points to PCIF1 and the downstream SCAP-cholesterol inflammatory output as potential entry points to expose immune vulnerability in poorly inflamed tumours. Any such strategy will need to consider tumour specificity, T cell-intrinsic PCIF1 functions and systemic cholesterol biology. More broadly, our findings suggest that tumour metabolism is not interpreted by the immune system in isolation, but is filtered through post-transcriptional regulatory gates that determine whether a metabolic programme remains cell autonomous or becomes immune visible.

## Methods

### Cell lines

HEK293T cells (CRL-3216), YUMM1.7 melanoma cells (CRL-3362) and Lewis lung carcinoma LLC1 cells (CRL-1642) were obtained from the American Type Culture Collection (ATCC). BP melanoma cells (*Braf^V600E^/Pten^-/-^)* were provided by Jennifer Wargo. HEK293T cells were used for lentiviral production, whereas YUMM1.7, LLC1 and BP cells were used as syngeneic murine tumour models. All cell lines were maintained in Dulbecco’s modified Eagle’s medium (DMEM; Gibco, C11995500BT) supplemented with 10% heat-inactivated fetal bovine serum (FBS; Lonsera, S711-001S). Cells were cultured at 37 °C in a humidified incubator containing 5% CO□ and were passaged before reaching full confluence. Cells were routinely monitored for morphology and growth characteristics and were tested for mycoplasma contamination using a Mycoplasma Detection Kit (SouthernBiotech, 13100-01). Only mycoplasma-negative cultures were used for experiments. Cell-line authentication was performed by short tandem repeat (STR) profiling. Cells were used within a limited number of passages after thawing, and independently generated frozen stocks were used to minimize culture-associated drift.

### In vivo tumour growth

All animal experiments were performed in accordance with protocols approved by institutional animal care and use committee of Sichuan University (No. 20210224152). Mice were housed under specific pathogen-free conditions with controlled temperature, humidity and a 12 h light-dark cycle, with free access to food and water. Six-week-old *C57BL/6J* mice (GemPharmatech, strain no. N000013) were used as immunocompetent syngeneic hosts. Six-week-old *Balb/cNj-Foxn1^nu^/Gpt* mice (GemPharmatech, strain no. D000521) were used as immunodeficient athymic hosts. Male mice were used for LLC1 and YUMM1.7 tumour studies, and female mice were used for BP melanoma tumour studies, unless otherwise indicated.

For subcutaneous tumour implantation, control cells expressing a non-targeting shRNA (shNTC) or Pcif1-depleted cells expressing shRNAs targeting Pcif1 (shPcif1) were collected during exponential growth, washed and prepared as single-cell suspensions in sterile 1×PBS. Cell viability was assessed by trypan blue exclusion, and only preparations with viability greater than 90% were used for implantation. LLC1 cells were implanted subcutaneously at 5 × 10^5^ cells per mouse into the right flank of male *C57BL/6J* or *Balb/cNj-Foxn1^nu^/Gpt* mice. YUMM1.7 cells were implanted subcutaneously at 1 × 10^6^ cells per mouse into the right flank of male *C57BL/6J* or *Balb/cNj-Foxn1^nu^/Gpt* mice. BP melanoma cells were implanted subcutaneously at 1 × 10^6^ cells per mouse into the right flank of female *C57BL/6J* or *Balb/cNj-Foxn1^nu^/Gpt* mice.

Tumour growth was monitored three times per week using digital callipers. Tumour volume was calculated using the formula: volume = length × width^2^ / 2, where length represents the longest tumour diameter and width represents the perpendicular diameter. Mice were euthanized when tumours reached the humane endpoint defined in the approved animal protocol, when tumour ulceration or impaired welfare was observed, or at the experimental endpoint. Tumour volume did not exceed 2000 mm^3^ in any experiment. Mice were assigned to experimental groups according to randomization method; for example, random allocation after tumour implantation or after tumour establishment, and investigators were blinded to group allocation during tumour measurement and analysis.

### Immune checkpoint blockade treatment in vivo

For immune checkpoint blockade experiments, LLC1 or YUMM1.7 tumour cells expressing shNTC or shPcif1 were implanted subcutaneously as described above. For combined checkpoint blockade in the LLC1 model, mice bearing shNTC or shPcif1 LLC1 tumours were randomized on day 6 after tumour inoculation and treated intraperitoneally with anti-PD1 antibody (clone 29F.1A12; Bio X Cell, BE0273; RRID: AB_2687796) together with anti-CTLA4 antibody (clone 9D9; Bio X Cell, BE0164; RRID: AB_10949609), or with the corresponding IgG isotype control antibodies (Bio X Cell, BE0089; RRID: AB_1107769). Antibodies were administered in 100 μl InVivoPure dilution buffer on days 6, 9 and 12 after tumour implantation. Anti-PD1 was given at 200 μg per mouse; anti-CTLA4 was given at 200 μg per mouse. For YUMM1.7 immune checkpoint blockade experiments, 1 × 10^6^ shNTC or shPcif1 YUMM1.7 cells were implanted subcutaneously per mouse. Mice were randomized on day 8 after tumour inoculation and treated intraperitoneally with anti-PD1 antibody or IgG isotype control at 200 μg per mouse in 100 μl InVivoPure dilution buffer on days 8, 11 and 14. Tumour growth was monitored three times per week by calliper measurement, and tumour volume was calculated as described above. Tumours did not exceed 2000 mm^3^.

### In vivo CD8□ T cell depletion

For CD8□ T cell depletion experiments, mice were randomized to receive anti-CD8a antibody or isotype control. Anti-CD8a antibody (clone 2.43; Bio X Cell, BE0061; RRID: AB_1125541) or rat IgG2b isotype control antibody (Bio X Cell, BE0090; RRID: AB_1107780) was administered intraperitoneally. A loading dose of 500 μg antibody in 100 μl InVivoPure dilution buffer (Bio X Cell, IP0070 or IP0065) was given one day before tumour implantation. After tumour implantation, maintenance doses of 250 μg antibody in 100 μl buffer were administered intraperitoneally twice weekly. For LLC1 depletion studies, 5 × 10^5^ shNTC or shPcif1 LLC1 cells were implanted subcutaneously per mouse, and anti-CD8a or isotype control treatment was continued twice weekly for two weeks after tumour inoculation. Tumour growth was measured three times per week using callipers, and tumour volume was calculated as (length × width^2^) / 2. Tumours did not exceed 2000 mm^3^. The efficiency of CD8□ T-cell depletion was assessed by immunofluorescence in tumour tissue.

### shRNA-mediated Pcif1 knockdown

Stable Pcif1 knockdown was achieved using lentiviral shRNA transduction. Lentiviral particles were produced in HEK293T cells using a calcium phosphate transfection method. Briefly, HEK293T cells were seeded in 10 cm tissue culture dishes to reach approximately 60–80% confluence at the time of transfection. Before transfection, the culture medium was replaced with fresh complete DMEM. Cells were co-transfected with the lentiviral shRNA plasmid (Horizon discovery, RMM3981-201829329 (Clone ID: TRCN0000111781) and RMM3981-201822737 (Clone ID: TRCN0000111783)) targeting mouse Pcif1 or a non-targeting control shRNA plasmid, together with the packaging plasmids psPAX2 (Addgene, #12260) and pMD2.G (Addgene, #12259), using calcium phosphate precipitation. After 6–8 h, the transfection medium was replaced with fresh complete medium. Virus-containing supernatants were collected 36 h after transfection, clarified by centrifugation at 2,500 rpm for 10 min at 4 °C, and filtered through a 0.45 μm filter before use. For generation of stable knockdown cell lines, YUMM1.7, LLC1 and BP cells were seeded to reach 60–80% confluence at the time of transduction. Cells were incubated with virus-containing supernatant supplemented with polybrene (10 μg.ml□¹; Solarbio, #H8761) for 6 h. The viral supernatant was then replaced with fresh complete medium containing puromycin (1.5 μg ml□¹; Selleck, S7417). Cells were maintained under puromycin selection for two weeks to generate stable pooled populations. Knockdown efficiency was validated by immunoblotting before cells were used for in vitro assays or in vivo tumour implantation. Stable cell populations were further maintained without puromycin selection and used within a limited number of passages after validation.

The shRNA target sequences used in this study were as follows:

mouse Pcif1

shRNA_781, 5’-AAACCCAGGGCTGCCTTGGAAAAG-3’

shRNA_783, 5’-TAATTTCACGAAACATGGATG-3’

non-targeting control

shRNA, 5’-CAACAAGATGAAGAGCACCAA-3’.

### Antibodies and related reagents

The primary antibodies used in this study were as follows: mouse anti-β-actin (ZSGB-BIO, TA-09; RRID: AB_2636897; 1:3,000 for immunoblotting), mouse anti-β-tubulin (ZSGB-BIO, TA-10; RRID: AB_3095964; 1:20,000 for immunoblotting), rabbit anti-SREBP2 (Thermo Fisher Scientific, PA1-338; RRID: AB_2194237; 1:1,000 for immunoblotting), rabbit anti-PCIF1 (Abcam, ab205016; RRID: AB_2753142; 1:1,000 for immunoblotting), mouse anti-IκBα (L35A5; Cell Signaling Technology, 4814T; RRID: AB_390781; 1:1,000 for immunoblotting), rabbit anti-NFκB p65 (D14E12; Cell Signaling Technology, 8242; RRID: AB_10859369; 1:1,000 for immunoblotting), rabbit anti-phospho-NFκB p65 Ser536 (Cell

Signaling Technology, 3033; RRID: AB_331284; 1:1,000 for immunoblotting), rabbit anti-SCAP (Proteintech, 12266-1-AP; RRID: AB_10697683; 1:1,000 for immunoblotting), rabbit anti- TBK1(Cell Signaling Technology, 3013S, RRID:AB_2199749; 1:1,000 for immunoblotting), rabbit anti-phospho-TBK1(Cell Signaling Technology, 5483S, RRID:AB_10693472; 1:1,000 for immunoblotting), rabbit anti-STAT1(Cell Signaling Technology, 14994S, RRID:AB_2737027; 1:1,000 for immunoblotting), rabbit anti-phospho-STAT1(Cell Signaling Technology, 9167S, RRID:AB_561284; 1:1,000 for immunoblotting), rabbit anti-IRF3(Cell Signaling Technology, 4302S, RRID:AB_1904036; 1:1,000 for immunoblotting), rabbit anti-phospho-IRF3(Cell Signaling Technology, 4947S, RRID:AB_823547; 1:1,000 for immunoblotting), rabbit anti-STING (Cell Signaling Technology, 13647T, RRID:AB_2732796; 1:1,000 for immunoblotting). Horseradish peroxidase-conjugated secondary antibodies used for immunoblotting were goat anti-rabbit IgG-HRP (ZSGB-BIO, ZB2301; RRID: AB_2747412; 1:10,000) and □goat anti-mouse IgG-HRP (ZSGB-BIO, ZB2305; RRID: AB_2747415; 1:10,000).

The following chemical reagents were used where indicated: cycloheximide (MedChemExpress, HY-12320), U18666A (MedChemExpress, HY-107433), UT-59 (DC Chemicals, DC74218) and methyl-β-cyclodextrin (MβCD; MedChemExpress, HY-101461). Cycloheximide was prepared as a 100 mg ml□¹ stock solution in DMSO and stored at −20 °C. U18666A and UT-59 were prepared as stock solutions in DMSO at 10 mM and stored at −20 °C. MβCD was prepared fresh in sterile PBS or culture medium at 500 mM before use, unless otherwise indicated. For all inhibitor or chemical perturbation experiments, vehicle-treated cells were used as controls, and the final concentration of DMSO was kept constant between treatment groups. Working concentrations and treatment durations were as follows: cycloheximide, 100 μg ml□¹ for 5 min; U18666A, 7.5 µg/ml for 12 h; UT-59, 3 μM for 3 h; and MβCD, 1 mM for 6 h.

### In vitro cell proliferation assay

In vitro proliferation of control and Pcif1-depleted tumour cells was assessed using the Cell Counting Kit-8 assay (CCK-8; Beyotime, C0038), which measures WST-8 reduction as a surrogate of viable cell number and metabolic activity. Cells were collected during logarithmic growth, dissociated into single-cell suspensions and counted using haemocytometer. Equal numbers of cells were seeded into 96-well tissue-culture plates at 5 × 10^3^ cells per well in 100 μl complete medium. For each cell line and condition, cells were plated in n = 3 technical replicate wells per time point. Medium-only wells containing CCK-8 reagent were included as background controls. Cell proliferation was measured at 0, 24, 48, 72 and 96 h after seeding. At each time point, 5 μl CCK-8 reagent was added directly to each well and cells were incubated for 2 h at 37 °C in a humidified incubator with 5% CO□. Absorbance at 450 nm was then measured using a microplate reader (Biotek Epoch II).

Background absorbance from medium-only wells was subtracted from all readings. Relative proliferation was calculated by normalizing the background-corrected absorbance at each time point to the corresponding 0 h value for the same cell line and condition, unless otherwise indicated. Experiments were performed using independently generated shNTC and shPcif1 cell populations and repeated in n = 3 independent biological experiments. For all comparisons, control and Pcif1-depleted cells were seeded at the same density and assayed in parallel on the same plate.

### Western blot analysis

Cells were harvested at approximately 70–80% confluence unless otherwise indicated. After two washes with ice-cold PBS, cells were lysed on ice in RIPA lysis buffer (Cell Signaling Technology, 9806S) supplemented immediately before use with DTT (1 mM, Sangon Biotech, A620058), PMSF (1 mM; Solarbio, P0100), 1x protease inhibitor cocktail (Solarbio, P6730) and 1x phosphatase inhibitor cocktail (Solarbio, P1260). Cells were scraped into lysis buffer, transferred to microcentrifuge tubes and incubated on ice for 30 min with intermittent vortexing. Lysates were clarified by centrifugation at 17,000g for 10 min at 4 °C, and the supernatant was collected as the soluble whole-cell extract.

Protein concentration was determined using a bicinchoninic acid assay (BCA; Solarbio, PC0020) according to the manufacturer’s instructions. Equal amounts of protein (40 μg) were mixed with 4× Laemmli sample buffer (Bio-Rad, 1610747) and denatured at 95 °C for 10 min. Proteins were separated by SDS–PAGE using 8% and 10% polyacrylamide gels according to the expected molecular weight of the target proteins. In particular, 8% gels were used for high-molecular-weight proteins including SCAP and SREBP2, whereas 10% gels were used for other targets. Proteins were transferred to nitrocellulose membranes (0.45 μm; Merck Millipore, HATF00010) using wet transfer in transfer buffer (25mM Tris, 192mM Glycine, 0.1%SDS, 20% ethanol) at 100 voltage for 1h 10min at 4 °C. After transfer, membranes were blocked for 1 h at room temperature in 5% non-fat milk in PBST. For detection of phosphorylated proteins, including phospho-NFκB p65 Ser536, membranes were blocked in 5% bovine serum albumin (BSA) in PBST to preserve phospho-epitope detection. Membranes were then incubated overnight at 4 °C with the indicated primary antibodies diluted in the corresponding blocking buffer. Primary antibodies and dilutions are listed in the Antibodies and reagents section.

After primary-antibody incubation, membranes were washed three times for 10 min each in PBST and incubated for 1 h at room temperature with HRP-conjugated secondary antibodies diluted in blocking buffer. The secondary antibodies used were HRP-conjugated goat anti-mouse rabbit IgG (ZSGB-BIO, ZB2301; RRID: AB_2747412; 1:10,000) and HRP-conjugated goat anti-mouse IgG (ZSGB-BIO, ZB2305; RRID: AB_2747415; 1:10,000). Membranes were washed three times for 10 min each in PBST, and signals were detected using an enhanced chemiluminescence substrate (4A Biotech, 4AW011). Chemiluminescent images were acquired using a ChemiDoc MP imaging system (Bio-Rad). Band intensities were quantified using Image Lab software (Bio-Rad). Immunoblots shown are representative of n = 3 independent biological experiments with similar results.

### Tumour cell conditioned medium cytokine array

Tumour-cell secreted inflammatory proteins were profiled using conditioned medium from control and Pcif1-depleted LLC1 cells. Cells were seeded at 5 × 10^5^ cells per 10 cm dish in complete DMEM and allowed to adhere overnight. The next day, cells were washed twice with sterile PBS to remove residual serum-derived proteins and incubated in serum-free DMEM for 6 h at 37 °C in 5% CO□. Conditioned medium was collected, centrifuged at 500g for 10 min at 4 °C to remove detached cells and debris, and stored at −80 °C until analysis. In parallel, cells from each dish were collected for protein quantification to allow normalization of cytokine signals to cell number or total cellular protein when required. Secreted cytokines and inflammatory mediators were screened using the Proteome Profiler Mouse XL Cytokine Array Kit (R&D Systems, ARY028), which detects 111 mouse cytokines and related proteins. For each sample, 500 μl conditioned medium was applied to the array membrane according to the manufacturer’s instructions. Membranes were incubated with conditioned medium and the supplied detection-antibody cocktail, followed by streptavidin–HRP incubation and chemiluminescent detection. Membrane images were acquired using a ChemiDoc MP imaging system (Bio-rad) under non-saturating exposure conditions. Array images were quantified using Fiji/ImageJ v2.1. For each analyte, the signal intensity of duplicate spots was measured after local background subtraction and averaged. Signals were normalized to the internal positive-reference spots on the same membrane. Normalized cytokine levels were then expressed relative to the corresponding shNTC control condition. Cytokines with signals below background or close to the negative-control spots were considered not detected and were excluded from fold-change interpretation. Data shown represent one screening experiment with n = 3 replicates. Candidate cytokines identified by the array, including CXCL10, were further validated by ELISA in independent conditioned-medium samples.

### Tumour tissue preparation for flow cytometry

Fresh tumour tissues were collected from tumour-bearing mice at the indicated experimental endpoints and processed immediately for flow cytometric analysis. Tumours were minced into small fragments with sterile scissors and enzymatically dissociated using the Tumour Dissociation Kit, mouse (Miltenyi Biotec, 130-096-730) according to the manufacturer’s instructions. Briefly, tumour fragments were incubated in the supplied enzyme mixture in DMEM at 37 °C for 10 min with intermittent agitation or using a gentleMACS dissociator programme. The resulting cell suspension was passed through a 40 μm cell strainer, and the strainer was washed with cold PBS containing 0.1% BSA. Cells were centrifuged at 300 g for 5 min at 4 °C, resuspended in staining buffer and counted. Red blood cells were lysed using RBC lysis buffer (Invitrogen, 00-4333-57). Cell viability was assessed using trypan blue, and equal numbers of cells were used for antibody staining.

For surface staining, single cell suspensions were first incubated with FcR Blocking Reagent (Miltenyi Biotec, 130-092-575; 1:50) for 15 min at 4 °C in PBS containing 0.1% BSA. Cells were then stained for 30 min at 4 °C in the dark with the indicated fluorophore-conjugated antibodies. Dead cells were excluded using the LIVE/DEAD Fixable Dead Cell Stain Kit (Thermo Fisher Scientific Invitrogen, L34966) according to the manufacturer’s instructions.

The antibodies used for flow cytometry were as follows: anti-mouse I-A/I-E (clone 2G9; BD Biosciences, 743870; RRID: AB_2741821; 1:100), anti-mouse Ly6C (clone HK1.4; BioLegend, 128049; RRID: AB_2800630; 1:100), anti-mouse Ly6G (clone 1A8; BD Biosciences, 740953; RRID: AB_2740578; 1:100), anti-mouse CD103 (clone 2E7; Miltenyi Biotec, 130-118-681; RRID: AB_2751551; 1:100), anti-mouse PD-1 (clone 29F.1A12; BioLegend, 135205; RRID: AB_1877232; 1:100), anti-mouse F4/80 (clone QA17A29; BioLegend, 157307; RRID: AB_2832550; 1:100), anti-mouse CD8a (clone 53-6.7; BD Biosciences, 564983; RRID: AB_2739032; 1:100), anti-mouse CD4 (clone RM4-5; BioLegend, 100525; RRID: AB_312726; 1:100), anti-mouse CD45 (clone 30-F11; BioLegend, 103130; RRID: AB_893339; 1:100), anti-mouse CD11c (clone N418; BioLegend, 117333; RRID: AB_11204262; 1:100) and anti-mouse NK1.1 (clone PK136; BioLegend, 108745; RRID: AB_2563286; 1:100). For intracellular Foxp3 staining, surface stained cells were fixed and permeabilized using 0.1% Triton X100 for 5 min, followed by staining with anti-mouse Foxp3 antibody (clone R16-715; BD Biosciences, 567373; RRID: AB_2916572; 1:100). Foxp3 staining was included only in experiments in which regulatory T-cell populations were analysed.

Flow cytometric acquisition was performed on a BD LSRFortessa cytometer using FACSDiva software v8. Compensation was performed using single bead stained compensation controls, and fluorescence-minus-one controls were used where required to define gates. Data were analysed using FlowJo software (Tree Star). The general gating strategy was as follows: single cells were selected by forward- and side-scatter parameters, dead cells were excluded by viability dye, and immune cells were identified as live CD45□ cells. CD8□ T cells were gated as live CD45□CD8a□ cells, and intratumoural CD8a□CD103□ T cells were quantified within the CD8□ T cell compartment. Tumour-associated macrophages were quantified as live CD45□F4/80□ cells, with additional myeloid markers including CD11c, I-A/I-E, Ly6C and Ly6G used to resolve myeloid subsets where indicated. NK cells were gated as live CD45□NK1.1□ cells. For experiments involving cell sorting, cells were sorted on a BD FACSAria II using the same staining and viability-exclusion strategy.

### Filipin III flow cytometery analysis

Free cholesterol was measured by Filipin III staining followed by flow cytometry. Cells were collected at the indicated time points, washed twice with ice-cold PBS and fixed in 4% paraformaldehyde in PBS for 30 min at room temperature. After fixation, cells were washed twice with PBS to remove residual paraformaldehyde and resuspended in freshly prepared Filipin III working solution (Sigma-Merck, SAE0088) diluted in PBS according to the manufacturer’s instructions. Cells were incubated for 1 h at 4 °C in the dark. After staining, excess Filipin III was removed by two washes with PBS, and cells were resuspended in PBS for immediate flow-cytometric analysis. Filipin III fluorescence was detected using UV excitation, with emission collected in the Pacific Blue-compatible channel (355-nm laser, 450-nm emission filter). Unstained fixed cells were used to define background fluorescence, and all samples within the same experiment were acquired using identical instrument settings. At least 10,000 single-cell events were collected per sample after exclusion of debris and cell aggregates by forward- and side-scatter parameters. Data were analysed using FlowJo 10.8.1 software (Becton Dickinson&Company). Filipin III signal was quantified as the percentage of Filipin III-high cells. Values were normalized to the shNTC control condition within each experiment where indicated.

### Polysome profiling

Polysome profiling was performed under conditions designed to preserve ribosome–mRNA complexes during cell harvest and fractionation. For each experimental condition, cells were seeded one day before the experiment at approximately 1 × 10^7^ cells per 15 cm dish, with at least two dishes prepared per condition. Cell number was adjusted according to cell size and growth rate so that cultures did not exceed 70% confluence at the time of harvest.

Sucrose gradients were prepared freshly on the day of the experiment. A 70% sucrose stock solution was prepared one day in advance in RNase-free Milli-Q water, dissolved completely with stirring, filtered through a 0.22 µm filter and stored at 4 °C. On the day of the experiment, 5% and 60% sucrose solutions were prepared from the 70% sucrose stock in gradient buffer. The final gradient buffer contained 20 mM HEPES pH 7.6, 100 mM KCl, 5 mM MgCl_2_, 100 ug ml^-1^ cycloheximide and RNase inhibitor. Linear 5–60% sucrose gradients were generated in SW41 ultracentrifuge tubes using a master gradient maker (Biocomp, Canada). Briefly, 5% sucrose solution was first added to the marked level of the tube, followed by underlaying of 60% sucrose solution from the bottom of the tube until the interface reached the same level. Tubes were sealed carefully to avoid bubble formation, and the gradient maker was run using the SW41 linear gradient programme to generate reproducible 5-60% gradients.

To arrest translation elongation and stabilize polysomes, cycloheximide was added directly to the culture medium at a final concentration of 100 μg ml^−1^, and cells were incubated for 5 min at 37 °C. Plates were then immediately transferred onto ice, and all subsequent steps were performed at 4 °C or on ice. Cells were washed twice with 5 ml ice-cold PBS containing 100 μg ml^−1^ cycloheximide and then scraped in 1 ml ice-cold PBS containing 100 μg ml^−1^ cycloheximide using a rubber cell scraper. Cell suspensions were collected in microcentrifuge tubes and centrifuged at 300g for 5 min at 4 °C. The supernatant was carefully removed. For each 15 cm dish, the cell pellet was resuspended in 450 μl freshly prepared hypotonic buffer containing 5 mM Tris-HCl pH 7.5, 2.5 mM MgCl_2_ and 1.5 mM KCl. Cycloheximide, DTT and RNase inhibitor were then added to final concentrations of 100 μg ml^−1^, 2 mM and 100 U ml^−1^, respectively. Lysates were briefly vortexed, after which Triton X-100 and sodium deoxycholate were added to final concentrations of 0.5% each. Samples were vortexed briefly again and incubated on ice for at least 30 min to allow complete lysis. Lysates were clarified by centrifugation at maximum speed for 5 min at 4 °C, and the supernatants were transferred to fresh tubes. When samples were not loaded immediately, clarified lysates were snap-frozen in liquid nitrogen and stored at −80 °C. Clarified lysates were quantified by absorbance, normalized across samples and loaded in equal ribosome equivalent amounts onto the sucrose gradients. Ten per cent of each normalized lysate was retained as input. Before loading, a volume of sucrose solution equivalent to the volume of lysate to be loaded was carefully removed from the top of each gradient. Samples were then gently layered onto the top of the gradients along the side of the tube to avoid perturbing the gradient. Tubes were balanced using hypotonic buffer. Gradients were centrifuged in an SW41 rotor using an Optima XPN ultracentrifuge at 36,000 rpm for 2 h 30 min at 4 °C. After centrifugation, gradients were fractionated using a Biocomp Gradient Station fractionator with continuous absorbance monitoring at 254 nm. Fractions corresponding to subpolysomal and polysomal ribosome populations were collected and snap-frozen in liquid nitrogen before storage at −80 °C or downstream analysis.

### Quantitative PCR analysis

Total RNA was extracted from cultured cells using TRIzol reagent (Invitrogen, Thermo Fisher Scientific, #15596018CN) according to the manufacturer’s instructions. Briefly, cells were lysed directly in TRIzol, and RNA was isolated by phase separation, precipitated with isopropanol, washed with 75% ethanol and resuspended in RNase-free water. RNA concentration and purity were determined using a NanoDrop microvolume spectrophotometer (Thermo Fisher Scientific), and equal amounts of RNA were used for reverse transcription across all samples.

Complementary DNA was synthesized using the PrimeScript RT Reagent Kit (Takara, #RR047A) according to the manufacturer’s instructions. For each sample, the same amount of total RNA was reverse-transcribed in parallel, and the resulting cDNA was diluted to an appropriate working concentration before quantitative PCR. Quantitative PCR was performed using SYBR Green Supermix (Bio-Rad, #1708882) on a CFX96 real-time PCR detection system (Bio-Rad). Reactions were set up using gene-specific primers and run under standard SYBR Green amplification conditions, followed by melting-curve analysis to confirm the specificity of amplification. Each biological sample was analysed with n = 3 technical replicates. Quantification cycle values were obtained using the Bio-Rad CFX software, and genes with non-specific amplification or abnormal melting curves were excluded from analysis. Relative mRNA expression was calculated using the ΔΔCq method.

Target gene expression was first normalized to the reference genes HPRT1. Relative expression values were then calculated against the indicated control condition and are presented as fold change unless otherwise specified. Primer sequences used for quantitative PCR are as follows:

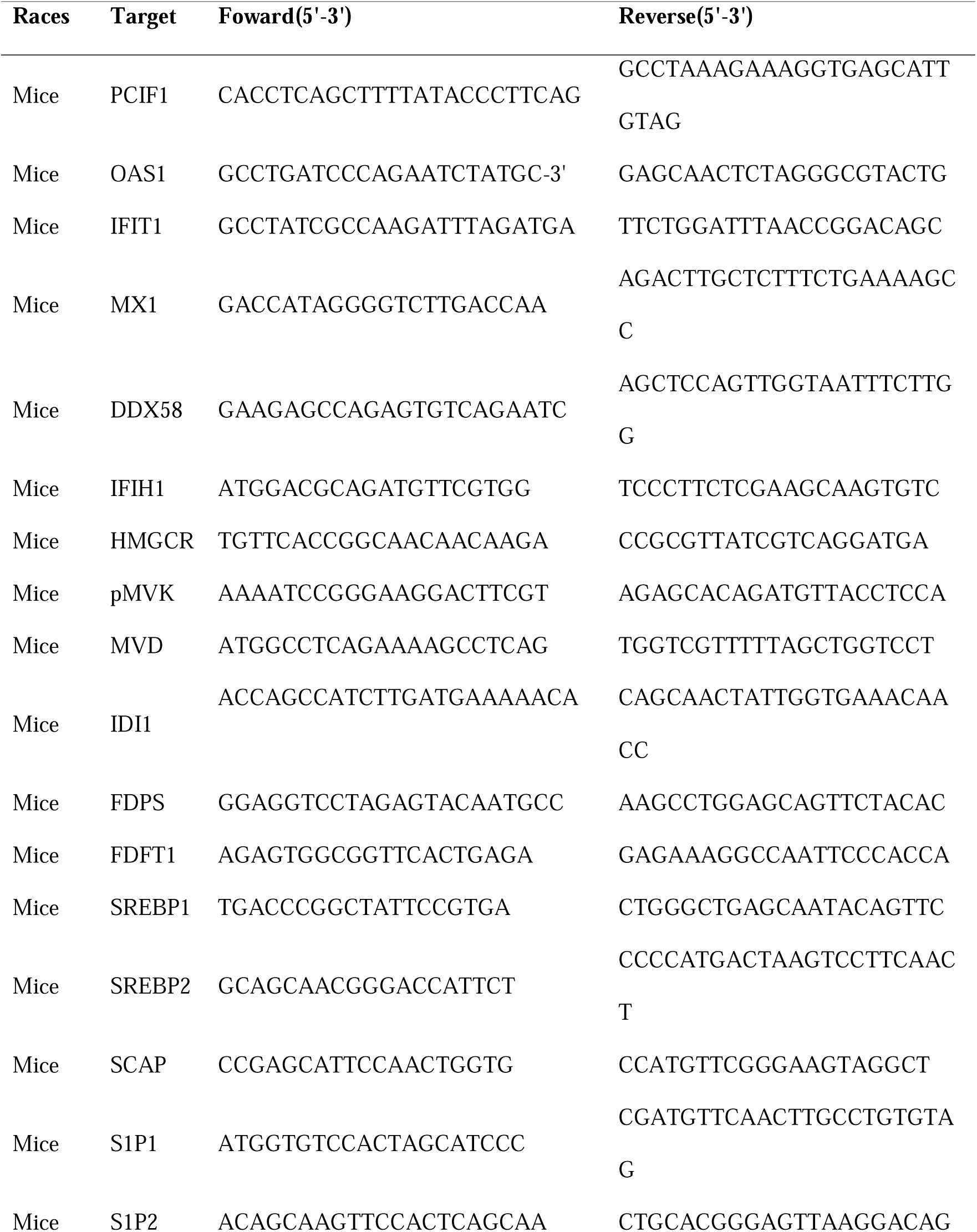
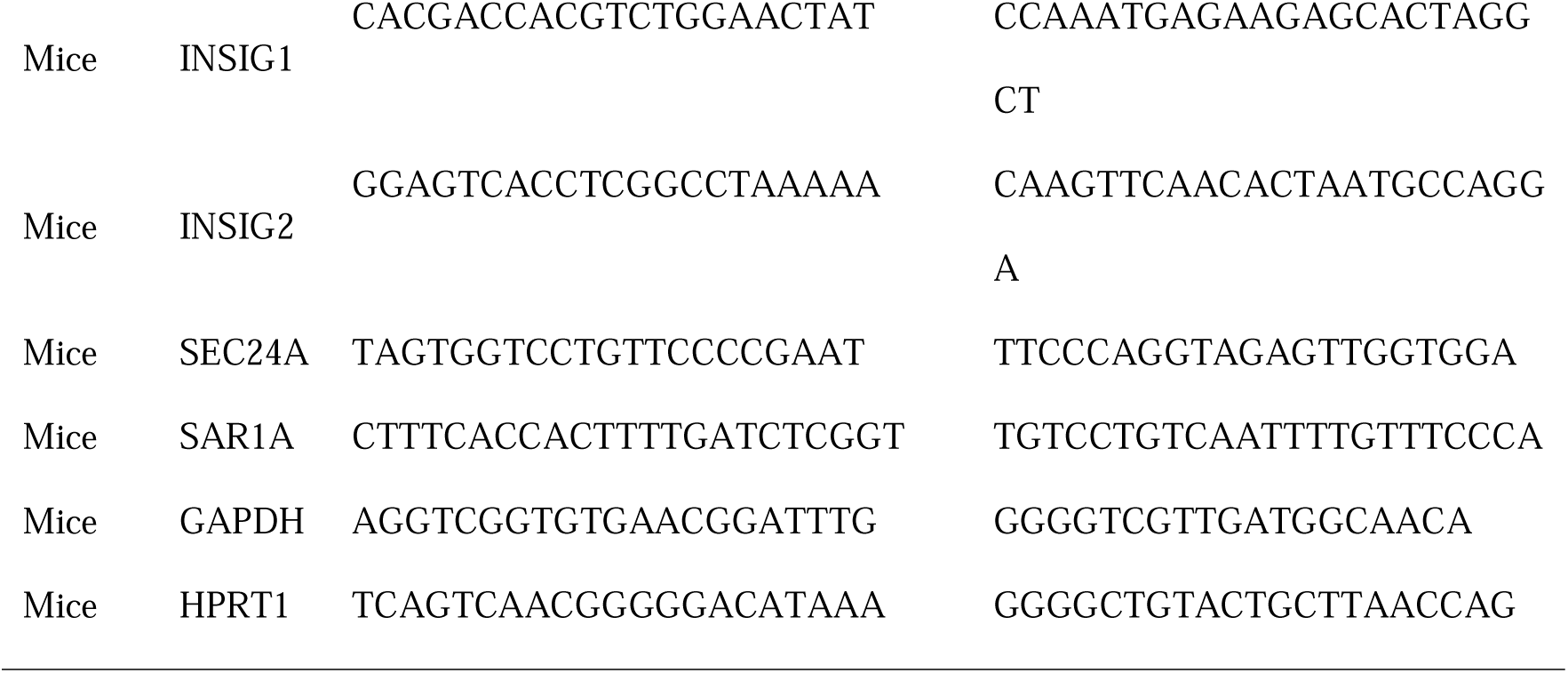

### CXCL10 ELISA analysis

To quantify secreted CXCL10, LLC1 sh_NTC and LLC1 sh_PCIF1 cells were seeded at 1 × 10^6^ cells per 60 mm dish and allowed to attach overnight. The next day, cells were washed once with pre-warmed PBS and cultured in fresh medium in the presence or absence of 1 mM methyl-β-cyclodextrin (MβCD; MedChemExpress, #HY-101461) for 6 h at 37 °C. At the end of treatment, conditioned media were collected and centrifuged at 2,500g for 10 min at 4 °C to remove detached cells and debris. The clarified supernatants were transferred to fresh tubes, aliquoted and stored at −80 °C until analysis. Repeated freeze–thaw cycles were avoided. CXCL10 concentrations in culture supernatants were measured using a mouse CXCL10 ELISA kit (Baipeng Bio, China, #bo04079) according to the manufacturer’s instructions. Briefly, standards and samples were brought to room temperature, added to the ELISA plate in four replicate wells per sample, and incubated with the capture and detection reagents supplied with the kit. After washing, substrate solution was added, the reaction was stopped according to the manufacturer’s protocol, and absorbance was measured using a microplate reader at 450 nm, with background correction at 630 nm. CXCL10 concentrations were calculated from a standard curve generated using recombinant CXCL10 standards and were expressed as pg ml^−1^. Samples with absorbance values outside the linear range of the standard curve were diluted and reanalysed. Where indicated, secreted CXCL10 levels were normalized to the number of viable cells measured in parallel at the time of supernatant collection.

### Immunofluorescence and immunohistochemistry

Tumour tissues were fixed in 4% paraformaldehyde (PFA, Beyotime Biotechnology) during overnight at room temperature, washed in PBS and transferred to 75% ethanol before paraffin embedding. Formalin-fixed, paraffin-embedded tissue blocks were sectioned at 4 μm using a microtome (LEICA, RM2245). Sections were de-paraffinized in xylene, rehydrated throughgraded ethanol solutions and rinsed in distilled water before antigen retrieval. Heat-induced epitope retrieval was performed in citrate antigen retrieval buffer, pH 6.0 (ZSGB-BIO, #ZLI9065), at 100 °C for 16 min. Sections were then allowed to cool to room temperature and washed in PBS.

For immunohistochemistry, endogenous peroxidase activity was quenched with hydrogen peroxide for 10 min at room temperature. Non-specific binding was blocked with goat serum (ZSGB-BIO, #ZLI9056) for 30 min at room temperature. Sections were incubated overnight at 4°C with the following primary antibodies: rabbit anti-PCIF1 (Abcam, #ab205016, RRID:AB_2753142, 1:200). After washing in PBS, sections were incubated with an anti-rabbit polymer secondary detection reagent (ZSGB-BIO, #PV-6000, RRID:AB_3675450) for 1 h at room temperature. Immunoreactivity was visualized using DAB substrate (Cell Signaling Technology, #11724S), and sections were counterstained with haematoxylin, dehydrated, cleared and mounted.

For multiplex immunofluorescence, staining was performed using the Opal Polaris 7-Color Automation IHC Kit (Akoya Biosciences, NEL871001KT) according to the manufacturer’s instructions. Briefly, paraffin sections were subjected to sequential rounds of antibody incubation, horseradish peroxidase-mediated Opal fluorophore deposition and heat-mediated antibody stripping. The following primary antibodies were used: rabbit anti-CD4 (Cell Signaling Technology, #25229S, RRID:AB_2798898, 1:200), anti-CD8 (Abcam, #ab217344, RRID:AB_2890649, 1:1,000), anti-CD19 (Abcam, #ab245235, RRID:AB_2895109, 1:500), anti-F4/80 (Cell Signaling Technology, #30325S, RRID:AB_2798990, 1:500) and anti-SREBP2 (Thermo Fisher Scientific, #PA1-338, RRID:AB_2194237, 1:400). Nuclei were counterstained with DAPI before mounting. Single-stained controls were used for spectral unmixing. Brightfield images of haematoxylin and eosin-stained and immunohistochemically stained sections were acquired using a VS200 whole-slide scanner (Olympus Life Science). Multiplex immunofluorescence slides were imaged using a Vectra Polaris imaging system (Akoya Biosciences), and multispectral images were unmixed using the corresponding single-stained controls. Tumour regions were manually delineated on whole-slide images by a trained pathologist who was blinded to experimental group allocation. Quantitative image analysis was performed using QuPath v0.4.3. Cells were detected on the basis of nuclear staining and expanded to approximate whole-cell boundaries. Positively stained cells were identified within pathologist-annotated tumour regions using marker-specific intensity thresholds that were kept constant across slides within the same staining batch. For multiplex immunofluorescence, marker-positive immune-cell populations were quantified as the proportion of CD4+, CD8+, CD19+ or F4/80+ cells among total nucleated cells within tumour regions, unless otherwise indicated.

### Bulk RNA-seq and pathway enrichment analysis

Total RNA was extracted from mouse-derived tumours and lymph nodes using TRIzol reagent (Invitrogen, Thermo Fisher Scientific, 15596018) according to the manufacturer’s instructions. RNA concentration and purity were assessed by spectrophotometry, and RNA integrity was evaluated using an Agilent 2100 Bioanalyzer. Samples with an RNA integrity number (RIN) ≥7.0 were used for library preparation. For each sample, polyadenylated RNA was enriched from total RNA using oligo(dT)-coupled magnetic beads. The purified mRNA was fragmented under elevated temperature in the presence of divalent cations, followed by first-strand cDNA synthesis using random hexamer primers and M-MuLV reverse transcriptase. RNA templates were subsequently removed by RNase H digestion, and second-strand cDNA was synthesized using DNA polymerase I and dNTPs. Double-stranded cDNA was subjected to end repair, 3′ adenylation and adaptor ligation. cDNA fragments of approximately 370–420 bp were selected using AMPure XP beads (Beckman Coulter), followed by PCR amplification and bead-based purification to generate the final sequencing libraries. Library concentration was initially measured using a Qubit 2.0 Fluorometer, and insert size distribution was assessed using an Agilent 2100 Bioanalyzer. Library molarity was further determined by quantitative PCR, and libraries with an effective concentration ≥2 nM were pooled according to target sequencing depth. Pooled libraries were sequenced on an Illumina NovaSeq 6000 platform to generate 150-bp paired-end reads. Raw image data were converted to base calls and FASTQ files using Illumina base-calling software. Raw reads were processed to remove adaptor sequences, reads containing ambiguous nucleotides and low-quality reads. Sequencing quality was assessed by calculating Q20 and Q30 scores, GC content and the overall clean-read yield. Only high-quality clean reads were used for downstream analyses. Clean paired-end reads were aligned to the mouse reference genome using HISAT2 (v2.0.5). The genome index was built from the reference genome together with the corresponding gene annotation file to enable splice-aware alignment. Gene-level read counts were obtained using featureCounts (v1.5.0-p3) based on uniquely mapped reads assigned to annotated genes. For visualization and descriptive analyses, transcript abundance was also normalized as fragments per kilobase of transcript per million mapped reads (FPKM), taking into account both sequencing depth and gene length. Differential expression analysis was performed on raw gene-level counts.

Differential gene expression between sh_NTC and sh_PCIF1 samples was analysed using the DESeq2 R package (v1.20.0), with three biologically independent samples per group. Genes with low read counts were filtered before statistical testing. DESeq2 normalization was applied to account for differences in sequencing depth across samples. Differential expression was estimated using a negative binomial generalized linear model, and P values were adjusted for multiple testing using the Benjamini–Hochberg method. Genes with an adjusted P value ≤0.05 and an absolute log2 fold change ≥1 were considered significantly differentially expressed.

Gene Ontology enrichment analysis was performed using the clusterProfiler R package (v3.8.1). Differentially expressed genes were analysed for enrichment of biological process, molecular function and cellular component terms, with correction for gene length bias where applicable. Enrichment P values were corrected for multiple comparisons, and GO terms with an adjusted P value <0.05 were considered significantly enriched.

### Polysome-associated RNA sequencing

Polysome profiling was performed as described above. After sucrose-gradient ultracentrifugation, gradients were fractionated with continuous absorbance monitoring at 254 nm, and ribosome-containing fractions were assigned on the basis of the absorbance profiles. For each sample, fractions corresponding to polysome-associated mRNAs were collected using the same fraction boundaries across all experimental conditions within the same experiment. Where indicated, equivalent polysome fractions were pooled before RNA extraction.

RNA was extracted from polysome fractions using TRIzol LS reagent (Invitrogen, Thermo Fisher Scientific) according to the manufacturer’s instructions, with reagent volumes adjusted to the volume of the collected sucrose fractions. RNA was precipitated, washed with 75% ethanol and resuspended in RNase-free water. RNA concentration and purity were measured using a NanoDrop microvolume spectrophotometer (Thermo Fisher Scientific). RNA integrity and quantity were further assessed before library preparation using an Agilent Bioanalyzer. Only RNA samples passing quality-control criteria for library construction were submitted for sequencing. RNA-seq libraries were generated by Novogene Bioinformatics Technology Co., Ltd. using the NEBNext Ultra RNA Library Prep Kit for Illumina (New England Biolabs, #E7530L) according to the manufacturer’s recommendations. Briefly, RNA samples were subjected to poly(A) mRNA enrichment, fragmented, reverse-transcribed and converted into indexed sequencing libraries after end repair, adaptor ligation and PCR amplification. Library quality was assessed by Qubit, and qualified libraries were pooled according to their effective concentrations. Pooled libraries were sequenced on an Illumina platform by Novogene Bioinformatics Technology Co., Ltd. using a paired-end 150-bp sequencing strategy. Raw sequencing reads were filtered to remove adaptor sequences, low-quality reads and reads containing ambiguous bases. Clean reads were aligned to the mouse mm10 reference genome using HISAT2. Gene-level read counts were generated using HTSeq based on Ensembl. Downstream normalization and differential expression analyses were performed in R using Xtail. Genes with low read counts were filtered before statistical analysis.

### Single cell RNA sequencing library preparation

Fresh tumour tissues were processed immediately after dissection to generate viable single-cell suspensions for single-cell RNA sequencing. Tumours were washed in cold PBS to remove blood and surface debris, transferred into GEXSCOPE Tissue Preservation Solution (Singleron Biotechnologies) and kept on ice until dissociation. For each experimental group, two tumour samples from different mice were undergoing tissue dissociation and library preparation independently. Tumour tissues were washed three times with Hank’s balanced salt solution and minced into small fragments using sterile blades. The minced tissues were incubated in Tissue Dissociation Solution (Singleron Biotechnologies) for 15 min at 37 °C with gentle agitation. After enzymatic dissociation, the cell suspension was passed through a 40 μm cell strainer to remove undigested tissue fragments and large debris. Cells were collected by centrifugation at 300g for 5 min, and the supernatant was discarded. The cell pellet was resuspended in PBS and subjected to red blood cell lysis using RBC lysis buffer. After lysis, cells were washed, centrifuged at 500g for 5 min and resuspended in PBS. Cell concentration and viability were determined by trypan blue exclusion using a haemocytometer. Only single cell suspensions with viability greater than 80% were used for subsequent library preparation. Single-cell suspensions were adjusted to a final concentration of 1 × 10^5^ cells ml^−1^ and loaded onto microfluidic chips using the GEXSCOPE Single-Cell RNA-seq Kit (Singleron Biotechnologies). Single-cell barcoding, reverse transcription and cDNA amplification were performed according to the manufacturer’s instructions. GEXSCOPE 3′ single-cell RNA-seq libraries were then constructed from amplified cDNA, indexed and pooled for sequencing. Library quality and concentration were assessed before sequencing according to the provider’s quality control workflow. Pooled libraries were sequenced on an Illumina NovaSeq platform using paired-end 150-bp reads.

### Single cell RNA sequencing data processing and analysis

Raw single-cell RNA-sequencing data were processed using the 10x Genomics Cell Ranger pipeline v7.1.0 with default parameters. Base-call files were demultiplexed into FASTQ files using cellranger mkfastq or the sequencing facility’s equivalent demultiplexing workflow.

Gene-expression libraries were processed using cellranger count. For each sample, sequencing reads were assigned to cell barcodes and unique molecular identifiers (UMIs) using the 10x barcode whitelist and the default Cell Ranger barcode-correction algorithm. Read 2 sequences were aligned to the mouse genome and transcriptome using the STAR aligner implemented in Cell Ranger. The reference used for alignment and gene annotation was the 10x Genomics mouse reference refdata-gex-mm10-2020-A. Reads confidently mapped to annotated exons and assigned to valid cell barcodes and UMIs were used for gene-level UMI counting. PCR duplicates were collapsed on the basis of cell barcode, UMI and gene assignment, and a gene-by-cell UMI count matrix was generated for each sample. Cell calling was performed using the default Cell Ranger algorithm. The filtered feature-barcode matrices, including matrix, barcode and feature files, were used as input for downstream analysis.

Downstream quality control, normalization, integration, clustering and visualization were performed using TrailMaker (Parse Bioscience). The Cell Ranger filtered feature-barcode matrices were imported into TrailMaker, and cell-level quality-control filtering was applied before integration. Cell-size distribution filtering was performed manually, with a minimum threshold of 1,000 transcripts per cell and a bin step of 200. Mitochondrial transcript content was filtered using a maximum mitochondrial percentage of 5%. The bin step for mitochondrial-content filtering was set to 0. The relationship between the number of detected genes and transcript counts was assessed using the automatic gene-number-versus-transcript-count filter with a linear fit, a prediction interval of 0.99992 and a P-value threshold of 0.000091. Putative doublets were removed using the automatic doublet filter with a probability threshold of 0.4 and a bin step of 0.02.

After quality control, data normalization and integration were performed in TrailMaker using the Seurat analysis module. Gene-expression values were normalized using the LogNormalize method. The 2,000 most highly variable genes were selected for downstream analysis. Ribosomal genes were excluded from the feature set used for integration and dimensionality reduction. Batch and sample effects were corrected using Harmony integration. Principal-component analysis was performed on the integrated expression matrix, and 30 principal components were retained for downstream analysis, accounting for 90.05% of the variation in the integrated dataset.

For visualization, cells were embedded using t-distributed stochastic neighbour embedding (t-SNE). t-SNE was performed with a perplexity of 30 and a learning rate of 978.1667. Graph-based clustering was performed using the Louvain algorithm at a resolution of 0.8. Clusters were annotated by combining canonical marker-gene expression, cluster-enriched genes and biological identity inferred from the tumour model. Major annotated cell populations included epithelial tumour cells, fibroblasts, T cells, NK cells, neutrophils, monocytes, tumour-associated macrophages, proliferating tumour-associated macrophages, cDC1, cDC2 and mature regulatory dendritic cells.

Cell-type annotation was supported by established marker genes. Immune cells were identified by Ptprc expression. T cells were annotated using Cd3d, Cd3e, Trac and Trbc2; cytotoxic lymphocyte states by Nkg7, Gzma, Ifng, Ccl5 and Xcl1; NK cells by cytotoxic lymphocyte genes in the absence of T-cell receptor markers; neutrophils by S100a8, S100a9, Csf3r, Hdc and G0s2; monocyte and macrophage populations by Cd14, Csf1r, Adgre1, C1qa, Folr2, Fcrls, Ccl7, Ccl12 and Arg1; dendritic-cell populations by Itgax, Flt3, Xcr1, Wdfy4, Cd209a, H2-Oa, Cd74, Ciita, Ccr7 and Ccl22; fibroblasts by Col1a1, Col1a2, Postn, Fbln2 and Serping1; and proliferating cell states by Mki67 and Cdca8. Epithelial tumour cells were annotated on the basis of tumour-associated epithelial markers and their separation from non-malignant immune and stromal populations.

For T-cell-state analysis, cells annotated as T cells were extracted from the integrated dataset and re-analysed separately using the same TrailMaker/Seurat workflow. T-cell subclusters were annotated on the basis of marker-gene expression. CD4 T cells were identified by Cd4 expression; cytotoxic CD8 T cells by Cd8a, Gzmb, Nkg7, Ifng, Ccl4, Ccl5 and Tnf; progenitor-like or memory-associated CD8 T-cell states by Tcf7, Il7r, Ccr7 and Sell; exhausted CD8 T-cell states by Tox, Pdcd1, Lag3, Havcr2 and Entpd1; interferon-stimulated CD8 T cells by Ifit1, Ifit2, Ifit3 and Isg15; γδ T cells by Trdc and Trgc1; and regulatory CD4 T-cell-associated features by Foxp3, Ctla4 and Tnfrsf4. For visualization, normalized gene expression was shown as feature plots, dot plots, heat maps or violin plots as indicated in the figure legends. For dot plots, dot size represents the percentage of cells expressing the indicated gene and colour represents scaled average expression. For heat maps and violin plots, expression values are shown as z-scored or log-normalized expression, as indicated. Cell-type proportions were calculated as the fraction of annotated cells within each sample or condition, or within the relevant parent compartment. Comparisons of selected marker-gene expression across shNTC, sh_781 and sh_783 groups were performed using the Kruskal–Wallis test unless otherwise indicated.

### InferCNV copy number variation analysis

Copy-number variation-like transcriptional profiles were inferred from single-cell RNA-sequencing data using inferCNV v1.13.0. The analysis was performed to support the identification of malignant epithelial tumour cells within the LLC1 tumour single-cell dataset and to distinguish tumour epithelial cells from non-malignant stromal and immune populations. The quality-controlled gene-by-cell UMI count matrix generated after Cell Ranger processing and downstream single-cell quality control was used as input. Cell annotations derived from the integrated TrailMaker/Seurat analysis were used to define query and reference populations. Cells annotated as epithelial cells were used as the query population. Non-malignant immune cells and fibroblasts were used as reference populations. These reference cells were selected because they represent host-derived diploid tumour microenvironment populations and showed no evidence of broad tumour-like epithelial transcriptional identity in the single-cell analysis. A cell annotation file was generated to assign each barcode to its corresponding cell type and experimental condition. A gene-order file was generated using the mouse mm10 genome annotation, with genes ordered according to chromosome and genomic coordinates. Genes not present in the expression matrix or without genomic-position information were excluded from the inferCNV input. The inferCNV object was generated using the filtered UMI count matrix, cell annotation file and mm10 gene-order file. inferCNV was run with immune and fibroblast populations specified as reference groups and with an expression cutoff of 0.1 to reduce noise from lowly expressed genes. Unless otherwise indicated, other parameters were kept at their default settings. Expression values from query epithelial cells were centred relative to the non-malignant reference populations, ordered along genomic coordinates and smoothed across neighbouring genes to infer large-scale chromosomal gains and losses from transcript abundance patterns. The resulting inferCNV profiles were visualized as chromosome-ordered heat maps. Cells were grouped by annotated cell type and experimental condition, and hierarchical clustering was used to display similarity among inferred copy-number profiles. Broad chromosome-scale deviations in epithelial cells relative to immune and fibroblast reference populations were interpreted as copy-number variation-like transcriptional patterns supporting malignant tumour-cell identity. Reference immune and fibroblast populations showed comparatively flat profiles, consistent with non-malignant diploid cells. The inferCNV analysis was used as orthogonal support for malignant epithelial-cell annotation and was not interpreted as direct genomic copy-number measurement.

### Lipidomic mass spectrometry sample preparation, data processing and analysis

Untargeted lipidomic profiling was performed on LLC1 cells expressing a non-targeting shRNA or shRNAs targeting Pcif1. Cells were cultured under matched conditions, washed twice with ice-cold PBS and harvested on ice. Cell pellets were snap-frozen in liquid nitrogen and stored at −80 °C until extraction. For each sample, lipids were extracted using an organic solvent-based extraction procedure. Briefly, frozen cell pellets were resuspended in ice-cold extraction solvent containing methanol and methyl tert-butyl ether (MTBE), vortexed and sonicated on ice to disrupt cells and extract lipids. Phase separation was induced by addition of water, followed by centrifugation at high speed at 4 °C. The upper organic phase containing lipids was collected, dried under nitrogen or vacuum centrifugation and reconstituted in isopropanol/acetonitrile-compatible injection solvent before liquid chromatography–mass spectrometry analysis. Pooled quality-control (QC) samples were generated by combining equal aliquots from all biological samples and were injected throughout the acquisition sequence to monitor instrument stability and analytical reproducibility. Solvent blanks were included to assess background and carryover signals. UHPLC–MS/MS analysis was performed by Novogene Co., Ltd. (Beijing, China) using a Vanquish UHPLC system coupled to an Orbitrap Q Exactive HF mass spectrometer (Thermo Fisher Scientific). Samples were separated on a Thermo Accucore C30 column (150 × 2.1 mm, 2.6 μm). The column temperature was maintained at 40 °C, and the flow rate was 0.35 ml min□¹. Mobile phase A consisted of acetonitrile/water (60:40, v/v) containing 10 mM ammonium acetate and 0.1% formic acid. Mobile phase B consisted of acetonitrile/isopropanol (10:90, v/v) containing 10 mM ammonium acetate and 0.1% formic acid. Lipids were separated using a 20-min gradient as follows: 30% B from 0 to 2 min, 43% B at 5 min, 55% B at 5.1 min, 70% B at 11 min, 99% B at 16 min and return to 30% B at 18.1 min, followed by column re-equilibration. Samples were analysed separately in positive- and negative-ion modes.

The Q Exactive HF mass spectrometer was operated in data-dependent acquisition mode. For positive-ion mode, the spray voltage was 3.5 kV, sheath gas was 40 psi, auxiliary gas was 10 arbitrary units, sweep gas was 0 arbitrary units, capillary temperature was 320 °C and auxiliary gas heater temperature was 350 °C. For negative-ion mode, the same settings were used except that the auxiliary gas was set to 7 arbitrary units. The S-lens RF level was 50. Full-scan MS spectra were acquired over an m/z range of 114-1,700 with an automatic gain control target of 3 × 10^6 and a maximum injection time of 100 ms. MS/MS spectra were acquired using stepped normalized collision energies of 22, 24 and 28 eV, with an isolation window of 1 m/z, an MS/MS automatic gain control target of 2 × 10^5 and a dynamic exclusion time of 6 s. Full MS and MS/MS Orbitrap resolution settings were set according to the standard Q Exactive HF lipidomics acquisition method used by the service provider and were kept identical for all samples.

Raw LC–MS/MS files were processed separately for positive- and negative-ion acquisition modes. Peak detection, retention-time alignment, peak integration, isotope/adduct grouping and lipid annotation were performed using Compound Discoverer 3 against an in-house and public lipid spectral library. Lipid features were retained when they had acceptable peak shape, signal-to-noise ratio and reproducible retention time across QC injections. Features detected in solvent blanks or dominated by blank signal were removed. Lipid annotations were assigned on the basis of accurate mass, retention time, isotope pattern and MS/MS fragmentation evidence where available. Features without sufficient annotation confidence were retained only for global feature-level analysis and were not used for lipid-class-level biological interpretation.

Peak intensities were normalized before downstream analysis to reduce technical variation across injections. Total ion current normalization was employed for this purpose. QC performance was assessed using pooled QC injections, total ion chromatograms and intensity distributions before and after normalization. Features with poor reproducibility in QC samples, defined as coefficient of variation greater than 30% unless otherwise indicated, were excluded from downstream analysis.

Multivariate analysis was performed separately for positive- and negative-ion datasets. Sparse partial least-squares discriminant analysis was used to visualize global lipidomic separation between control and Pcif1-depleted cells. Differential lipid features were identified by comparing Pcif1-depleted cells with control cells. For volcano plots and heat maps, features were considered significantly altered when they met both criteria: false-discovery rate (FDR) < 0.05 and |log2 fold change| > 1.5. FDR values were calculated by Benjamini-Hochberg correction of feature-level P values. Heat maps were generated using normalized, log2-transformed and z-scored feature intensities.

For lipid-class-level analysis, annotated lipid features were grouped according to lipid class. The median log2 fold change for each lipid class was calculated from all annotated features assigned to that class. Feature-level P values within each lipid class were combined using Fisher’s combined probability test to obtain a class-level P value. Class-level P values were then adjusted across lipid classes using the Benjamini–Hochberg procedure to calculate class-level FDR values. The number of significantly upregulated and downregulated features within each lipid class was determined using the same thresholds used for feature-level differential analysis, FDR < 0.05 and |log2 fold change| > 1.5. Lipid classes analysed included phosphatidylcholine (PC), phosphatidylethanolamine (PE), phosphatidylserine (PS), phosphatidylinositol (PI), phosphatidylglycerol (PG), phosphatidic acid (PA), cardiolipin (CL), sphingomyelin (SM), ceramide (Cer), glucosylceramide (GlcCer), bis(monoacylglycero)phosphate/hemibismonoacylglycerophosphate (BMP/HBMP), triacylglycerol (TAG), acylcarnitine (ACar) and fatty acid (FA) species, where detected. Lipid-class analyses were interpreted as evidence of lipidome remodelling rather than direct measurement of lipid metabolic flux.

### Quantification and statistical analysis

Statistical analyses were performed using GraphPad Prism v9.5.1. No statistical method was used to predetermine sample size. Sample sizes for cell-culture and animal experiments were chosen on the basis of previous experience with the corresponding assays and tumour models. Mice were randomly assigned to treatment groups where indicated. Investigators were not blinded to group allocation during in vitro or in vivo experiments. Technical replicates were not treated as independent biological replicates. The definition of n is provided in the corresponding figure legends and refers to biologically independent samples, mice, tumours, patient samples or independent experiments, as appropriate. Data are shown as mean ± s.e.m. unless otherwise indicated. All statistical tests were two-sided unless otherwise stated, and P < 0.05 was considered statistically significant. For comparisons between two independent groups, unpaired two-sided Student’s t-tests were used for approximately normally distributed data, and Mann–Whitney tests were used for non-parametric comparisons, including comparisons of PCIF1 expression between clinical response groups. For comparisons among three or more groups, one-way ANOVA was used where appropriate. Kruskal–Wallis tests were used for non-parametric comparisons among three groups, including selected single-cell marker-gene expression analyses across shNTC, sh_781 and sh_783 groups. Tumour-growth curves were analysed using two-way ANOVA. Survival curves from mouse experiments and patient cohorts were visualized using the Kaplan–Meier method and compared using log-rank tests. Hazard ratios and 95% confidence intervals for clinical survival analyses were obtained from the indicated survival-analysis platform. For qPCR, ELISA and flow-cytometry analyses, values were calculated from biologically independent samples before statistical testing. qPCR data were normalized to the indicated housekeeping gene. For single-cell RNA-seq analyses, cell-type proportions were calculated within the indicated parent population. Selected marker-gene expression comparisons among shNTC, sh_781 and sh_783 groups were performed using Kruskal–Wallis tests and interpreted as cell-state-associated differences. For gene-set enrichment analyses, enrichment statistics are reported as normalized enrichment scores and false-discovery-rate-adjusted q values. For lipidomics, differential lipid features were identified from normalized log-transformed intensities and were considered significant when both criteria were met: FDR < 0.05 and |log□ fold change| > 1.5. Feature-level P values were corrected using the Benjamini–Hochberg procedure. For lipid-class-level analysis, feature-level P values within each lipid class were combined using Fisher’s combined probability test, followed by Benjamini–Hochberg correction across lipid classes to calculate class-level FDR values. Associations between PCIF1 expression and cytolytic activity scores were assessed using Spearman correlation. Clinical survival and response analyses were retrospective and exploratory.

## Data Availability

RNA-seq data acquired in this study have been deposited in the Genome Sequence Archive of National Genomics Data Center, China National Center for Bioinformation, Chinese Academy of Sciences (https://ngdc.cncb.ac.cn/gsa). The accession number for RNA-seq data of LLC1 tumours and inguinal lymph nodes is GSA:CRA045545; the accession number for RNA-seq data of LLC1 and Yumm1.7 cell lines in polysome profiling is GSA:CRA045543; The single cell RNA-seq data of LLC1 tumours is accessible through GSA:CRA045548; Gene expression matrix of the single cell RNA-seq data reported in this paper have been deposited in the OMIX, China National Center for Bioinformation, Beijing Institute of Genomics, Chinese Academy of Sciences (https://ngdc.cncb.ac.cn/omix) with accession no.OMIX018282. The untargeted lipidomics data reported in this paper have been deposited in the OMIX, China National Center for Bioinformation, Beijing Institute of Genomics, Chinese Academy of Sciences (https://ngdc.cncb.ac.cn/omix) with accession no.OMIX018065.

Publicly available clinical and transcriptomic datasets were used for survival, immune checkpoint blockade response and cytolytic activity analyses. Survival analyses across lung adenocarcinoma, colon cancer, oestrogen receptor-positive breast cancer and serous ovarian cancer were performed using KM Plotter^44^ and the public datasets incorporated in the corresponding KM Plotter modules. The lung adenocarcinoma datasets included GSE14814, GSE19188, GSE29013, GSE30219, GSE31210, GSE3141, GSE31908, GSE37745, GSE50081 and GSE68465. The colon cancer datasets included GSE12945, GSE14333, GSE143985, GSE17538, GSE29621, GSE31595, GSE33114, GSE37892, GSE38832, GSE39582, GSE41258 and GSE92921. The oestrogen receptor-positive breast cancer datasets included GSE16716, GSE20711, GSE22093, GSE3494, GSE37946, GSE42568, GSE45255, GSE48390, GSE65194, GSE69031 and GSE7390. The serous ovarian cancer datasets included GSE14764, GSE15622, GSE26193, GSE30161, GSE32062, GSE63885, GSE65986, GSE9891 and TCGA ovarian cancer data. Public immune checkpoint blockade cohorts were used to assess the association between PCIF1 expression and treatment outcome. The glioblastoma anti-PD1 cohort was obtained from GSE121810. Melanoma anti-PD1 analyses used the Gide et al. 2019 cohort^45^, available through the European Nucleotide Archive under accession PRJEB23709, together with GSE78220, GSE91061 and the Liu et al. 2019 anti-PD1 melanoma cohort^46^. Melanoma anti-CTLA4 analyses used the Gide et al. 2019 cohort, GSE165278, the Liu et al. 2019 cohort and the Van Allen et al. 2015 cohort^47^. The Van Allen et al. 2015 melanoma anti-CTLA4 sequencing data are available through dbGaP under accession phs000452. Processed Gide et al. 2019 RNA-seq data are also available through the associated PredictIO/Zenodo resource with DOI 10.5281/zenodo.6968597. For cytolytic activity analyses, glioblastoma data were obtained from TCGA, lung adenocarcinoma data from GSE30219, breast cancer data from METABRIC and stage IV melanoma data from GSE22153. TCGA datasets were accessed through publicly available TCGA resources.

## Acknowledgements

S.S. acknowledges grant support from National Natural Science Foundation of China (grant no. 82473387) and “Rongpiao” innovative programme of Chengdu city, Sichuan, China. L. Liu acknowledges financial supports ZYGD18021 and ZYJC21002 from 1.3.5 Project for Disciplines of Excellence, West China Hospital, Sichuan University. C.R. thanks S. Bazin, O. and C. Courtin, Ensemble Contre le Mélanome, the Foundation Crédit Mutuel, Foundation Carrefour and the association Vaincre le Mélanome for their ongoing research funding support.

## Author Contributions

K.L., T.L. and Z.D. are co-first authors, contributed equally. K.L., T.L., S.S. designed the study. K.L., Z.D., S.S. interpreted the data and wrote the original draft of the manuscript. K.L., T.L., M.W., Y.W. performed RNA-seq and polysome profiling. L.K. performed in vivo experiment. D.Z., D.Y. performed bioinformatics analysis. S.R., N.B., E.E., J.Y.S., C.R. performed patient sample multiplex staining and developed image analysis pipeline. J.Y.S. performed histopathology analysis. L.L. provided administrative support and data resources, S.R., N.B., C.R.collected patient samples. S.S. supervised the overall study, provided strategic oversight, provided financial support and conceived the study.

