## Supplementary figures and images for "PCIF1 loss licenses antitumour immunity via cholesterol biosynthesis"

### Supplemental figure 1

##### Extended Data Figure 1

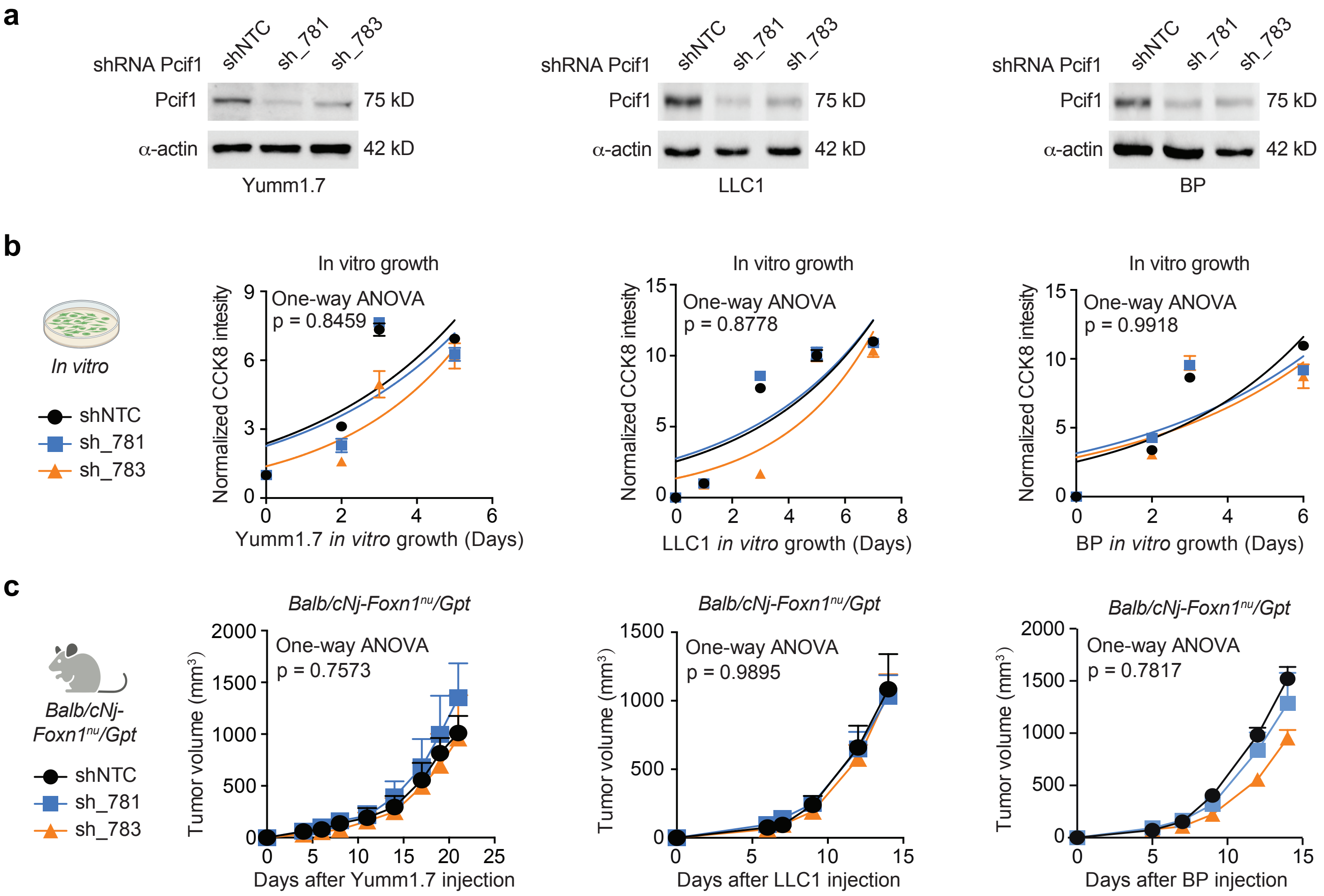

### Supplemental figure 2

**Extended Data Figure 2**

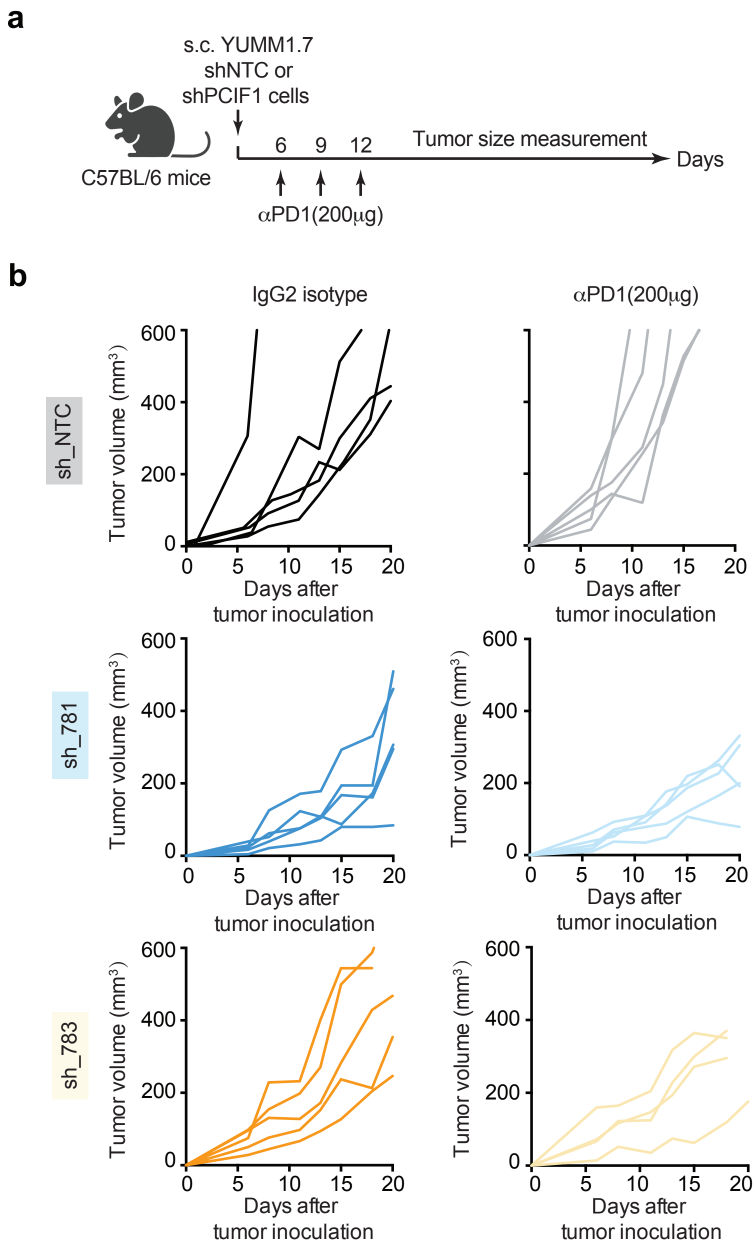

### Supplemental figure 3

#### Extended Data Figure 3

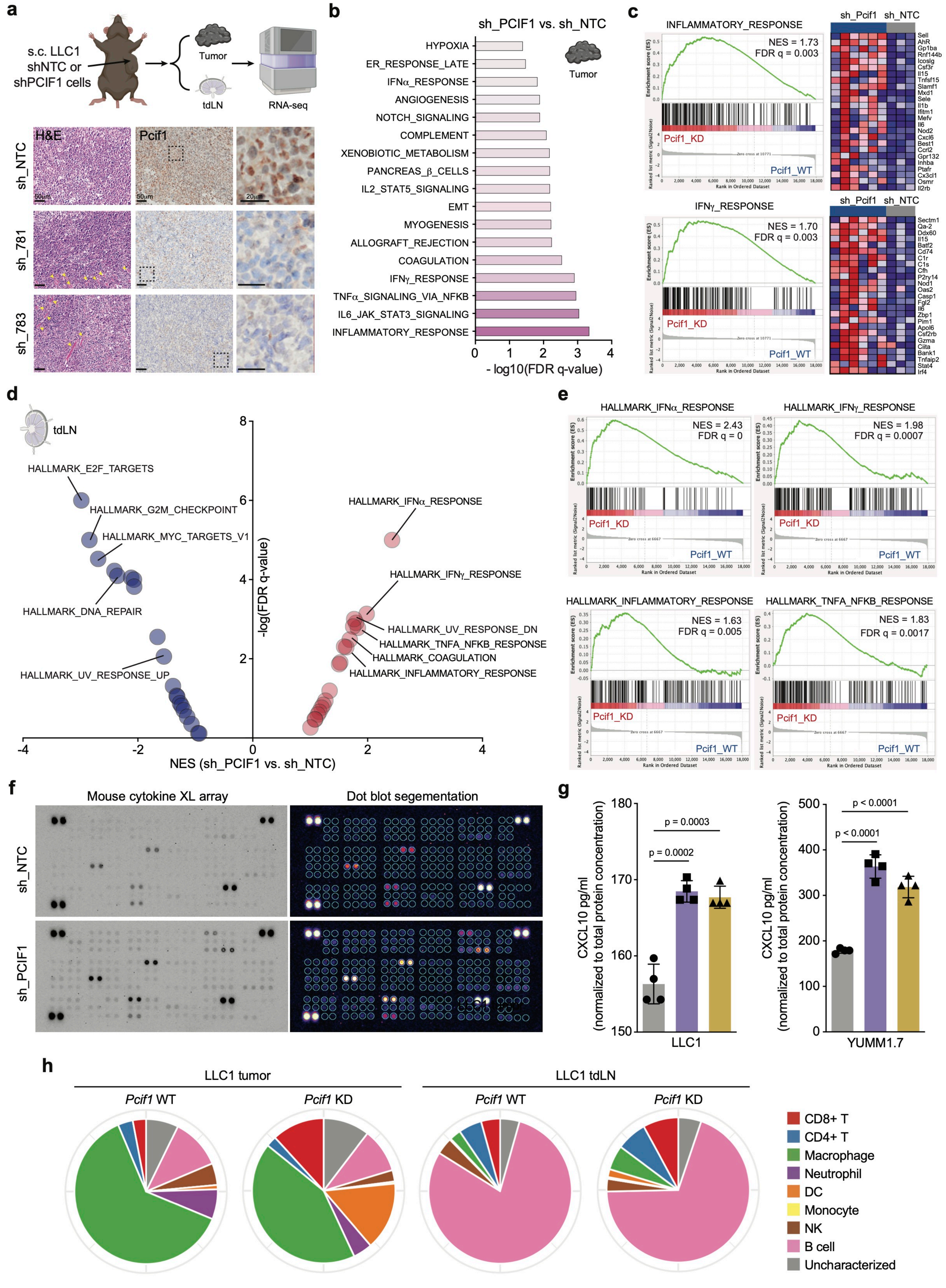

### Supplemental figure 4

#### Extended Data Figure 4

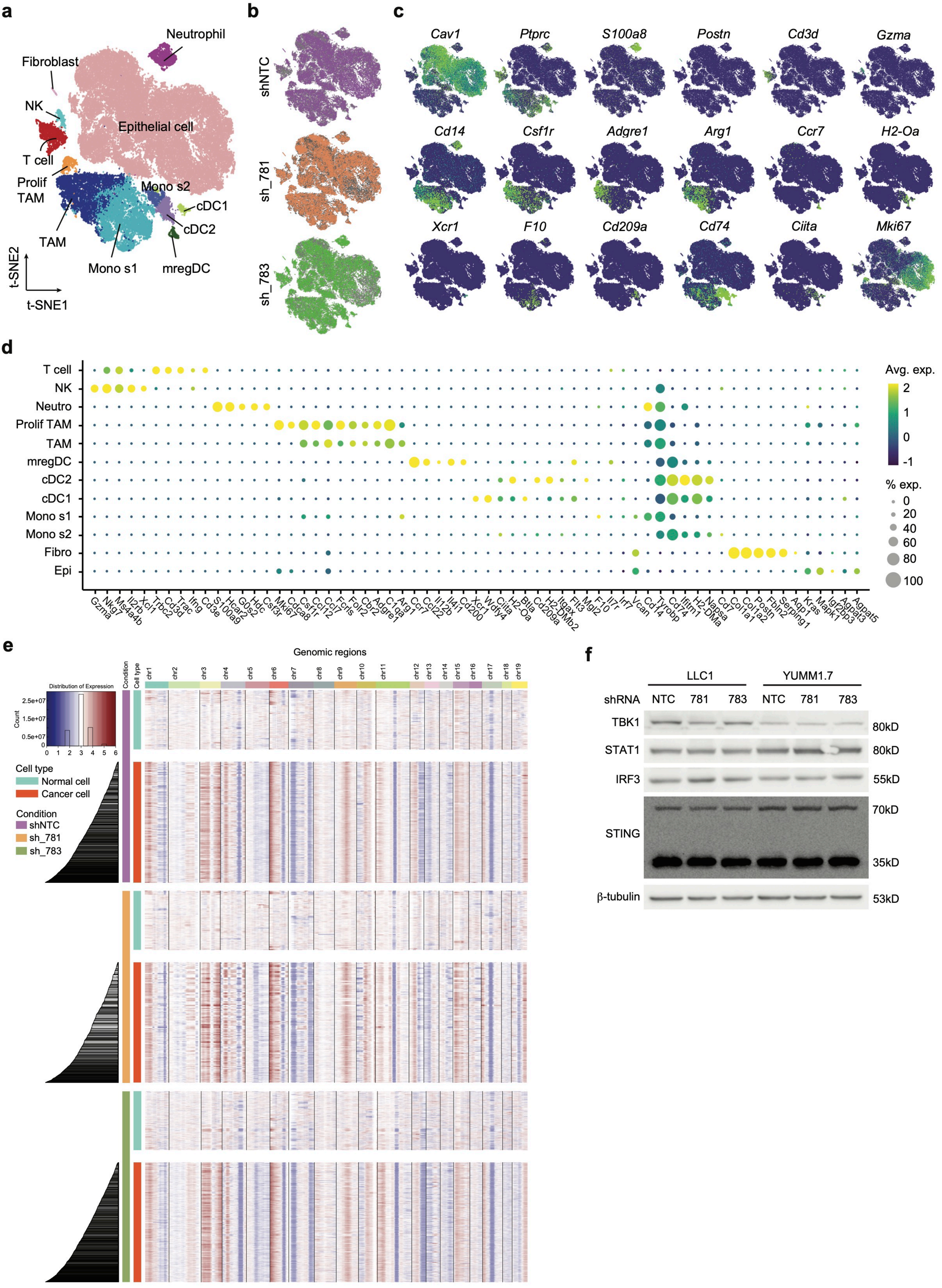

### Supplemental figure 5

**Extended Data Figure 5**

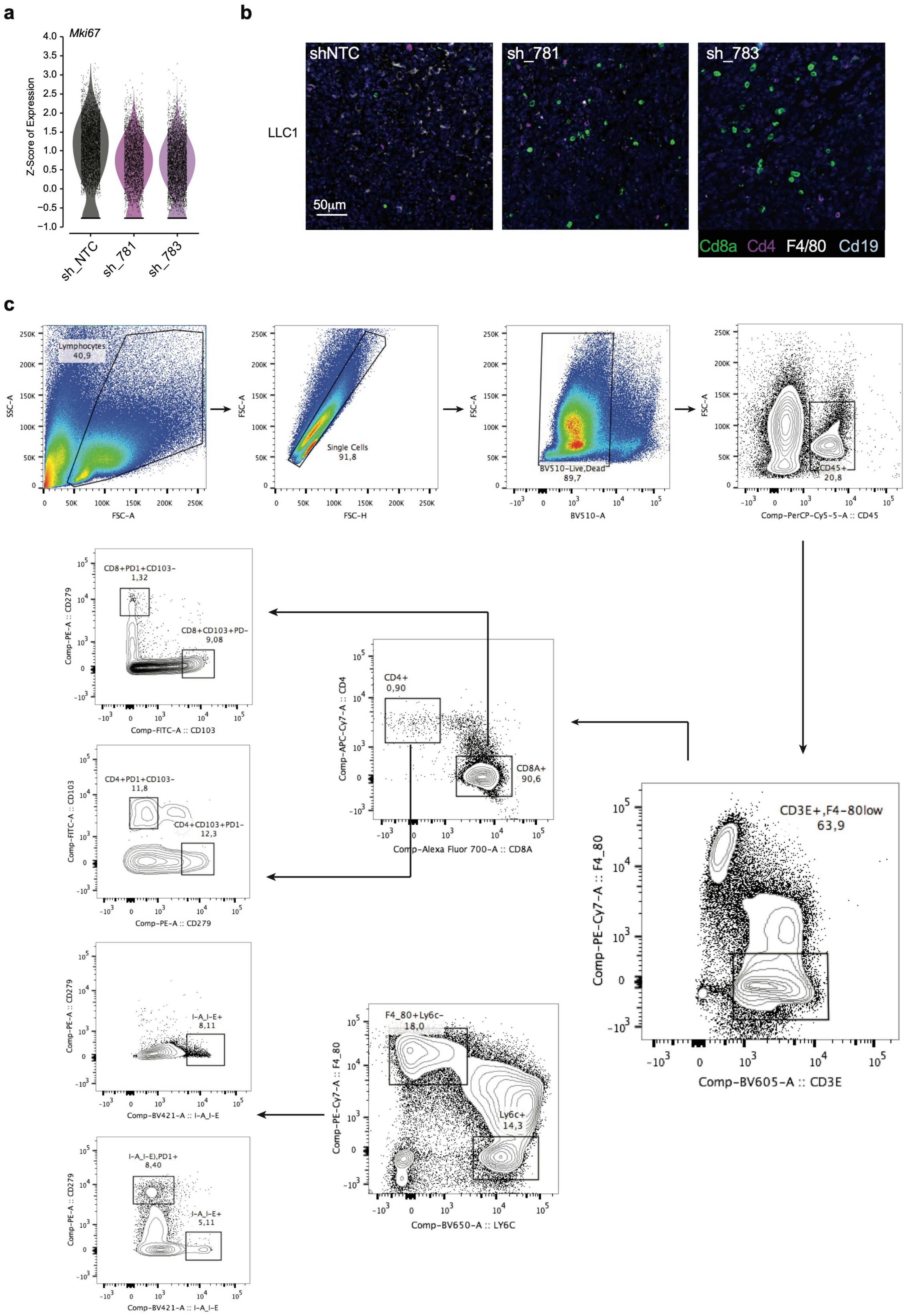

### Supplemental figure 6

##### Extended Data Figure 6

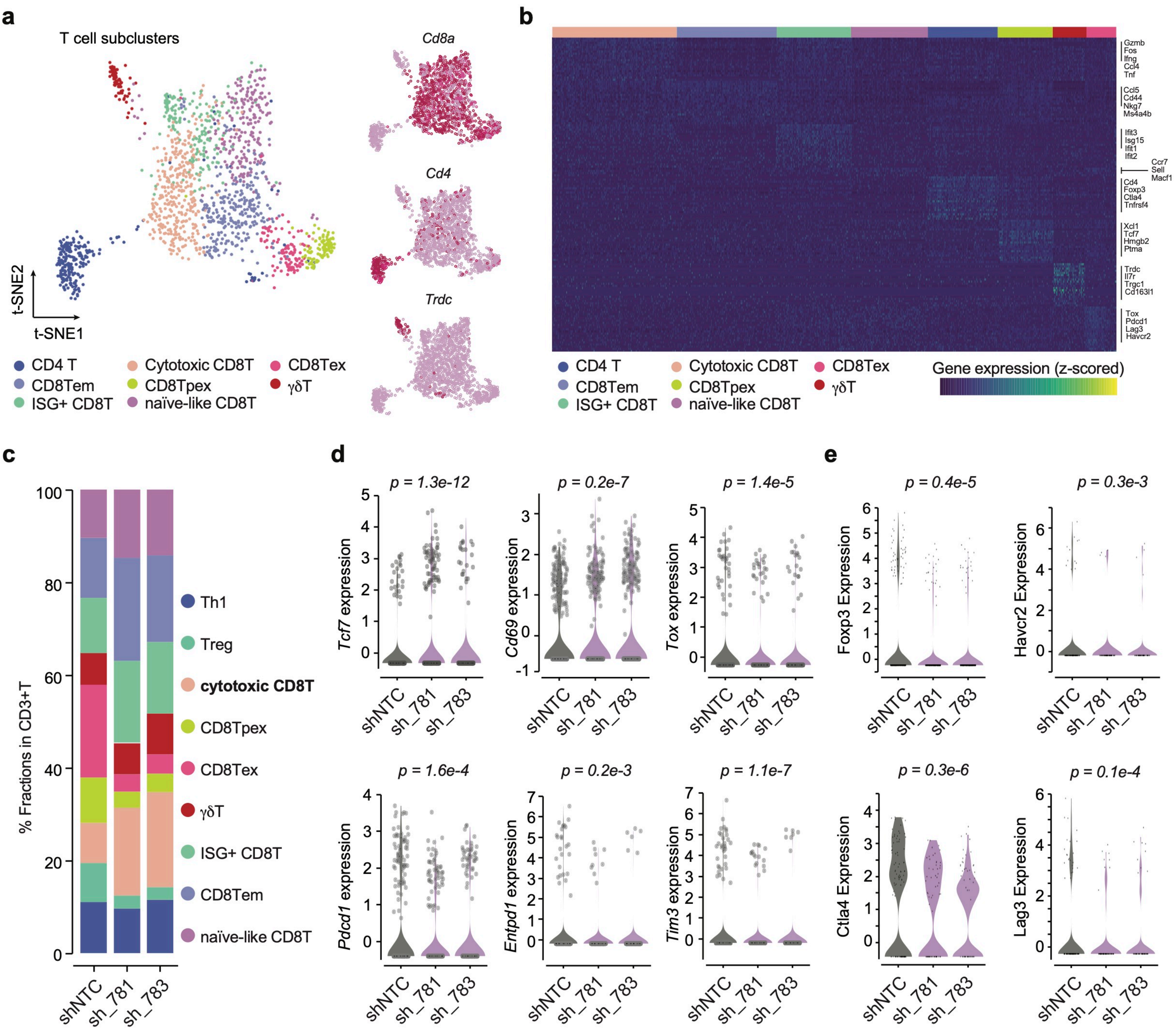

### Supplemental figure 7

#### Extended Data Figure 7

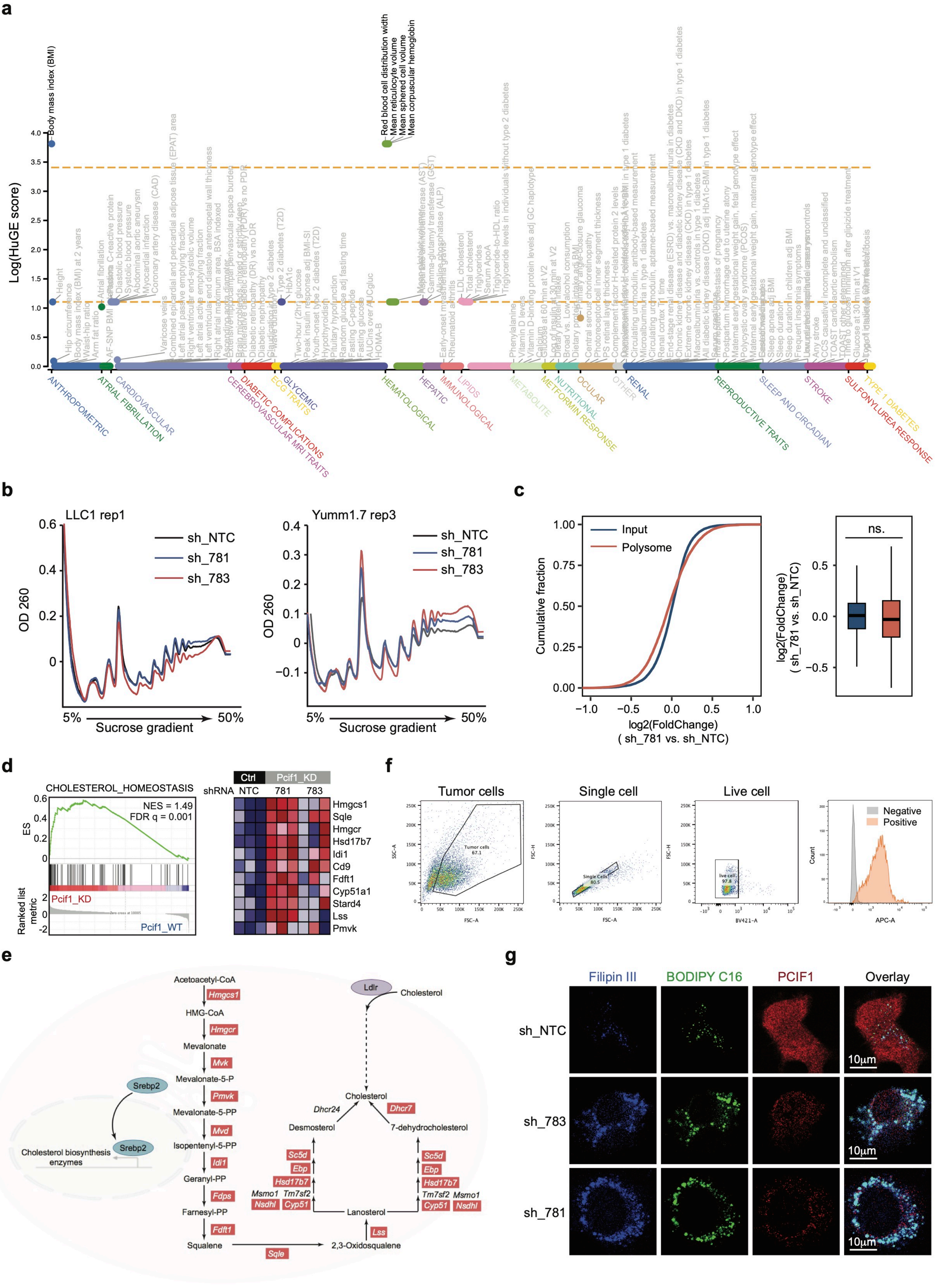

### Supplemental figure 8

##### Extended Data Figure 8

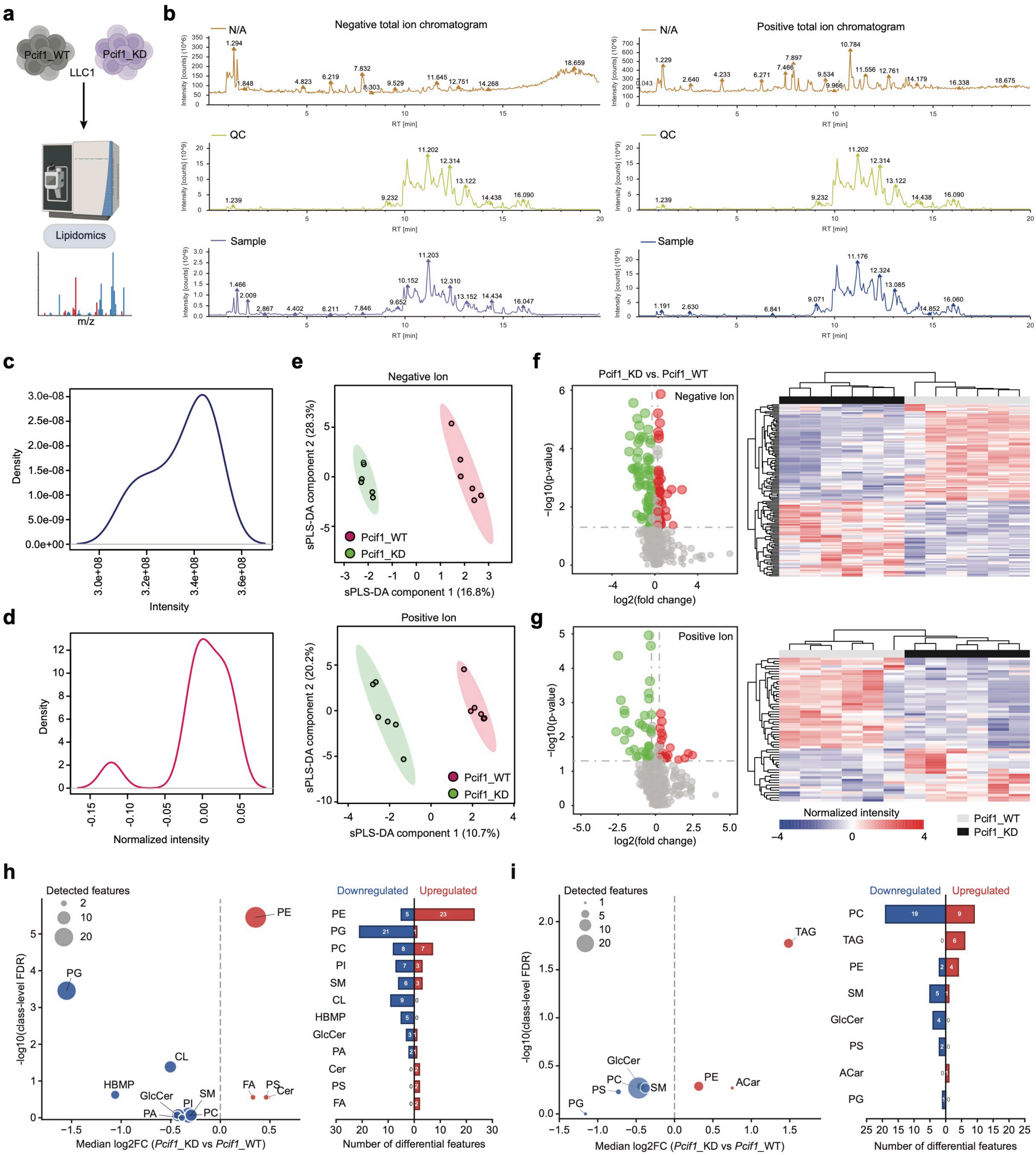

### Supplemental figure 9

##### Extended Data Figure 9

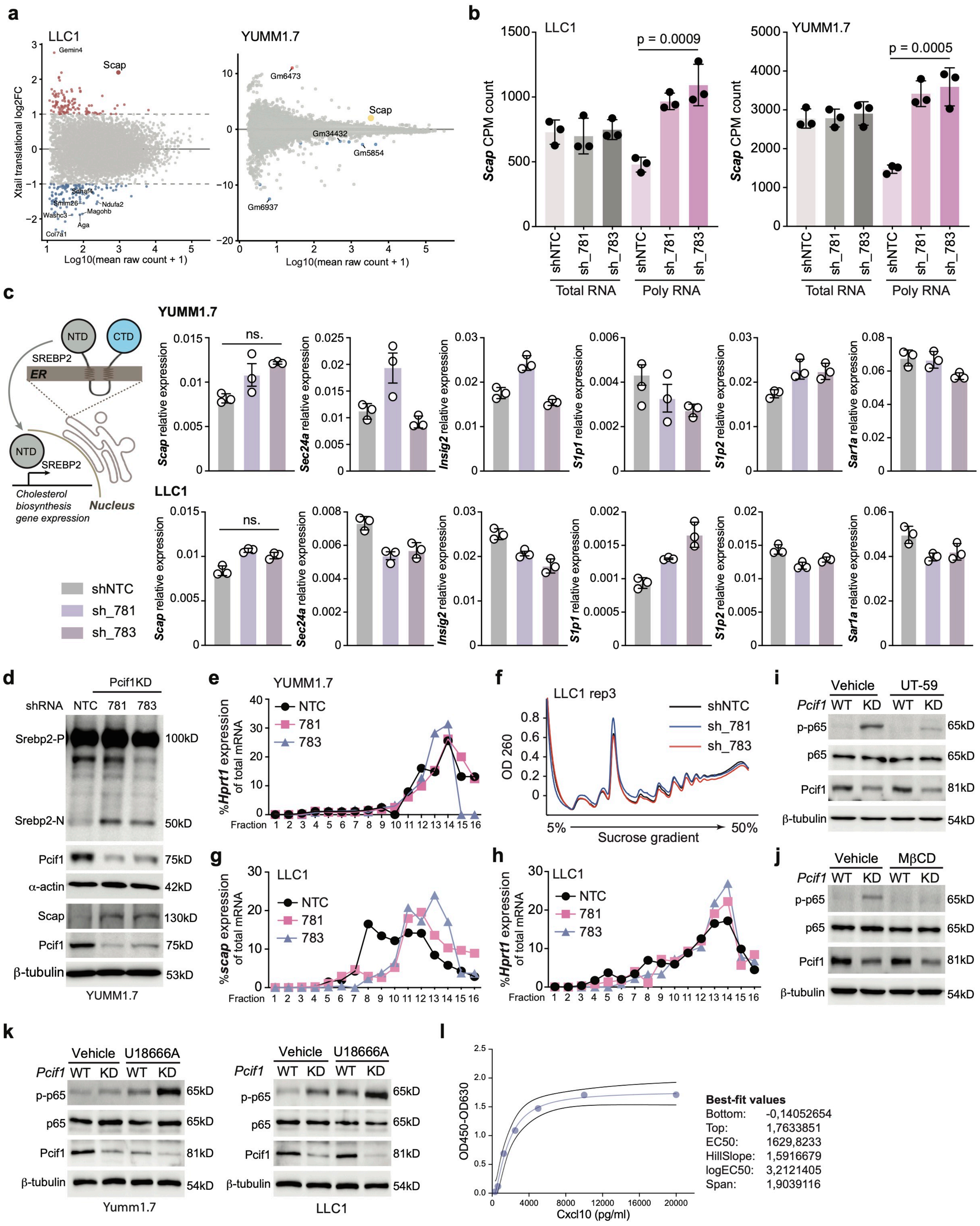

### Supplemental figure 10

#### Extended Data Figure 10

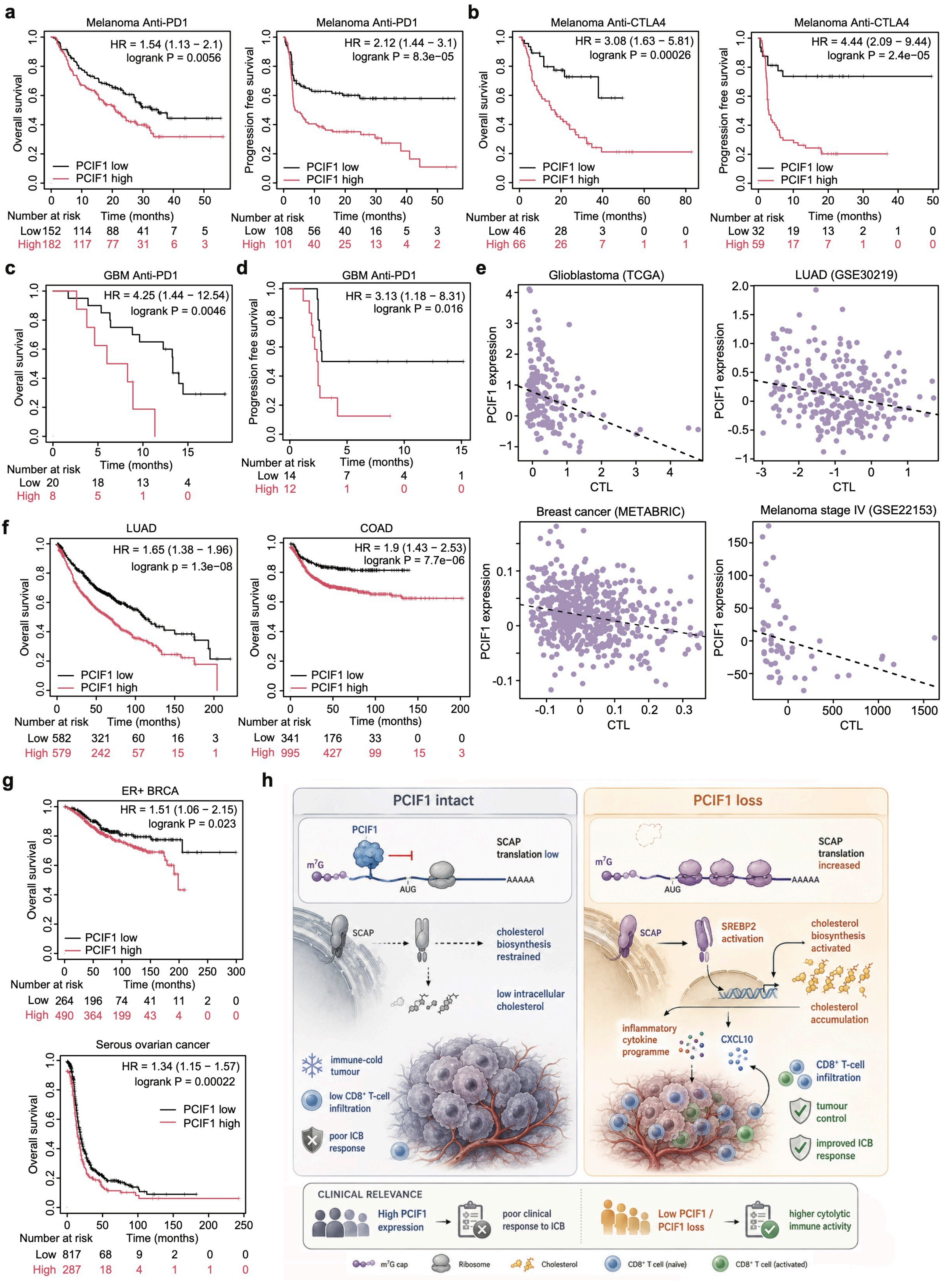
