## Supplemental Figure legend for "PCIF1 loss licenses antitumour immunity via cholesterol biosynthesis"

**Extended Data Figure Legend**

**Extended Data Figure 1. Pcif1 knockdown does not alter baseline tumour-cell growth in vitro or in immunodeficient mice.**

(**a**) Immunoblot analysis of PCIF1 expression in Yumm1.7, LLC1 and BP cells expressing a non-targeting control shRNA (shNTC) or two independent shRNAs targeting Pcif1 (sh_781 and sh_783). α-actin was used as a loading control. Molecular masses are indicated on the right. Blots are representative of n = 3 independent experiments.

(**b**) In vitro growth of Yumm1.7, LLC1 and BP cells expressing shNTC, sh_781 or sh_783, assessed by CCK8 assay at the indicated time points and shown as normalized CCK8 intensity. For statistical analysis, the area under the curve (AUC) was calculated for each biological replicate and compared among the three shRNA groups by one-way ANOVA. Exact P values are shown in the panels. Data are shown as mean ± s.e.m.; n = 3 biologically independent samples per group.

(**c**) Growth of subcutaneous Yumm1.7, LLC1 and BP tumours expressing shNTC, sh_781 or sh_783 in *Balb/cNj-Foxn1^nu^/Gpt* mice. Tumour volumes were measured at the indicated time points. For statistical analysis, the AUC of each tumour-growth curve was calculated for each mouse and compared among the three shRNA groups by one-way ANOVA. Exact P values are shown in the panels. Data are shown as mean ± s.e.m.; n = 5 mice per group.

**Extended Data Figure 2**. **Individual tumour-growth kinetics of Pcif1-knockdown Yumm1.7 tumours treated with PD-1 blockade.**

(**a**) Schematic of the in vivo treatment experiment. C57BL/6 mice were subcutaneously implanted with Yumm1.7 melanoma cells expressing a non-targeting control shRNA (shNTC) or one of two independent shRNAs targeting Pcif1 (sh_781 or sh_783). Mice were treated with IgG2 isotype control or anti-PD-1 antibody (200 μg per mouse, i.p.) on days 6, 9 and 12 after tumour inoculation. Tumour size was monitored at the indicated time points.

(**b**) Individual tumour-growth curves for Yumm1.7 shNTC, sh_781 and sh_783 tumours treated with IgG2 isotype control or anti-PD-1 antibody. Each line represents one mouse. Tumour volumes were measured longitudinally after tumour inoculation. For statistical analysis, the area under the curve (AUC) was calculated for each individual tumour growth curve and compared between IgG2- and anti-PD-1-treated mice within each shRNA group using one-way ANOVA test. n = 5 mice per group.

**Extended Data Figure 3**. **Pcif1 knockdown is associated with inflammatory and interferon-response programmes in LLC1 tumours and tumour-draining lymph nodes.**

(**a**) Experimental design for transcriptomic analysis of LLC1 tumours and tumour-draining lymph nodes (tdLNs). LLC1 cells expressing a non-targeting control shRNA (shNTC) or one of two independent shRNAs targeting Pcif1 (sh_781 or sh_783) were implanted subcutaneously into *C57BL/6J* mice. Tumours and matched tdLNs were collected on day 14 after tumour implantation for bulk RNA-seq analysis. Representative haematoxylin and eosin (H&E)-stained tumour sections and PCIF1 immunohistochemistry are shown for shNTC, sh_781 and sh_783 tumours. Boxed regions in low-magnification PCIF1-stained images are shown at higher magnification. Scale bars, 50 μm for low-magnification images and 20 μm for magnified images. Yellow arrows indicate the blood vessels.

(**b**) Hallmark pathway enrichment analysis of bulk RNA-seq data from LLC1 tumours comparing Pcif1-knockdown tumours with shNTC control tumours. Tumours generated from sh_781 and sh_783 cells were analysed collectively as the sh_PCIF1 group. Bars show −log_10-transformed FDR q values for the indicated enriched Hallmark pathways. Inflammatory-response, IL6-JAK-STAT3 signalling, TNFα signalling via NF-κB and IFNγ-response programmes were among the most enriched pathways in sh_PCIF1 tumours.

(**c**) Gene set enrichment analysis (GSEA) plots showing enrichment of the Hallmark inflammatory-response and IFNγ-response gene sets in sh_PCIF1 tumours compared with shNTC tumours. Pcif1_KD and Pcif1_WT indicate Pcif1-knockdown and control tumours, respectively. Normalized enrichment scores (NES) and FDR q values are indicated in the panels. Heat maps show relative expression of representative leading-edge genes from the indicated gene sets across individual tumour samples. Rows represent genes and columns represent tumour samples; expression values are shown as row-scaled z scores normalized expression values.

(**d**) Hallmark pathway enrichment analysis of bulk RNA-seq data from tdLNs associated with sh_PCIF1 versus shNTC LLC1 tumours. Each point represents one Hallmark gene set, plotted according to its NES on the x axis and −log_10-transformed FDR q value on the y axis. Positively enriched pathways in tdLNs from mice bearing sh_PCIF1 tumours are shown on the right, whereas negatively enriched pathways are shown on the left. Selected pathways are labelled, including IFNα response, IFNγ response, inflammatory response, TNFα signalling via NF-κB, allograft rejection and coagulation among positively enriched programmes, and E2F targets, G2M checkpoint, MYC targets and DNA repair among negatively enriched programmes.

(**e**) Representative GSEA enrichment plots for selected Hallmark pathways in tdLNs from mice bearing Pcif1-knockdown tumours compared with control tumours. IFNα-response, IFNγ-response, inflammatory-response and TNFα–NF-κB-response gene sets were enriched in tdLNs associated with Pcif1-knockdown tumours. NES and FDR q values are indicated in each plot.

(**f**) Mouse cytokine XL array analysis of conditioned medium from shNTC and sh_PCIF1 LLC1 tumour samples. Representative cytokine-array membranes are shown on the left, and the corresponding dot-blot segmentation used for signal quantification is shown on the right. Each pair of duplicate spots corresponds to an individual cytokine or control feature on the array.

(**g**) CXCL10 concentrations measured by ELISA in culture supernatants from LLC1 and Yumm1.7 cells expressing shNTC, sh_781 or sh_783. CXCL10 levels were normalized to total protein concentration. Data are shown as mean ± s.e.m.; P values were calculated using unpaired Mann-Whitney non-parametric analysis comparing each Pcif1 shRNA with shNTC. n = 4 biologically independent samples per group. Exact P values are shown in the panels.

(**h**) Estimated immune-cell composition of LLC1 tumours and matched tdLNs from Pcif1-wild-type and Pcif1-knockdown groups, inferred from bulk RNA-seq profiles using CIBERSORTx. Pie charts show the estimated relative proportions of the indicated cell populations, including CD8+ T cells, CD4+ T cells, macrophages, neutrophils, dendritic cells, monocytes, natural killer cells, B cells and uncharacterized cells. Pcif1 WT and Pcif1 KD denote control and Pcif1-knockdown groups, respectively. These values represent deconvolution-based estimated absolute cell fractions rather than direct cell counts. For **b-e**, pathway enrichment was performed using MSigDB Hallmark gene sets. For **b-e** and **h**, sh_781 and sh_783 samples were analysed jointly as the Pcif1-knockdown group. For **h**, immune-cell fractions were inferred using CIBERSORTx from bulk RNA-seq data and should therefore be interpreted as estimated proportions.

**Extended Data Figure 4**. **Single-cell annotation of LLC1 tumours and assessment of canonical innate immune signalling components after PCIF1 loss.**

(a) t-SNE visualization of single-cell RNA-seq profiles from LLC1 tumours generated from control or Pcif1-depleted tumour cells. Cells passing quality control were integrated and clustered, and major tumour, immune and stromal populations were annotated on the basis of canonical marker-gene expression. The annotated populations included epithelial tumour cells, fibroblasts, T cells, NK cells, neutrophils, monocytes, tumour-associated macrophages (TAMs), proliferating TAMs, conventional dendritic cell subsets (cDC1 and cDC2) and mature regulatory dendritic cells (mregDCs). Each dot represents one cell. The clusters were visualized with Trailmaker (Parse Biosciences).

(**b**) t-SNE projection of the same integrated single-cell dataset coloured by experimental condition. Cells were derived from LLC1 tumours expressing a non-targeting shRNA (shNTC) or two independent Pcif1 shRNAs (sh_781 and sh_783), showing the distribution of cells from each condition across the integrated cellular landscape. The sample-level clusters were visualized with Trailmaker (Parse Biosciences).

(**c**) Feature plots showing representative marker genes used to support cell-type annotation. Markers include genes associated with immune cells and lymphocytes (Ptprc, Cd3d and Gzma), neutrophils or inflammatory myeloid cells (S100a8), monocytes and macrophage-lineage cells (Cd14, Csf1r, Adgre1 and Arg1), dendritic-cell subsets and antigen-presentation states (Ccr7, H2-Oa, Xcr1, F10, Cd209a, Cd74 and Ciita), fibroblasts or stromal cells (Postn and Cav1), and proliferating cells (Mki67). Colour intensity indicates normalized expression of the indicated gene. The expression map was visualized with Trailmaker (Parse Biosciences).

(**d**) Dot plot showing expression of selected canonical marker genes across annotated cell populations. Dot size represents the percentage of cells expressing each gene, and colour represents scaled average expression. T-cell identity was supported by expression of Cd3d, Cd3e, Trac, Trbc2, Il2rb, Nkg7, Gzma, Xcl1 and Ifng; neutrophil identity by S100a9, Hcar2, G0s2, Hdc and Csf3r; proliferating TAMs by Mki67 and Cdca8; TAM and monocyte populations by Csf1r, Ccl12, Ccl7, Fcrls, Folr2, Cbr2, Adgre1, C1qa, Arg1, Vcan, Cd14, Tyrobp and Cd74; dendritic-cell subsets by Ccr7, Ccl22, Il12b, Il4i1, Cd200, Xcr1, Wdfy4, Ciita, H2-Oa, Btla, Cd209a, H2-DMb2, Itgax, Flt3, Mgl2 and F10; fibroblasts by Col1a1, Col1a2, Postn, Fbln2 and Serping1; and epithelial tumour cells by Kras, Mapk1, Igf2bp3, Agpat3 and Agpat5.

(**e**) inferCNV analysis of single-cell transcriptomes from LLC1 tumours. Non-malignant cell populations were used as reference cells to infer large-scale copy-number variation-like transcriptional patterns across genomic regions. Cells annotated as epithelial tumour cells showed broad chromosomal copy-number variation patterns that were not present in reference non-malignant populations, supporting their malignant identity. Rows represent individual cells ordered by cell type and experimental condition, and columns represent genomic regions ordered by chromosome.

(**f**) Immunoblot analysis of total TBK1, STAT1, IRF3 and STING protein abundance in LLC1 and YUMM1.7 cells expressing shNTC or two independent Pcif1 shRNAs (sh_781 and sh_783). β-tubulin was used as a loading control. In contrast to the increase in NFκB p65 phosphorylation shown in Fig. 2, PCIF1 loss did not cause a consistent increase in the total abundance of these canonical innate immune signalling components under the same experimental conditions. Blots are representative of n = 3 independent experiments with similar results.

**Extended Data Figure 5**. **Assessment of tumour-cell proliferation, immune-cell infiltration and flow-cytometric gating strategy in Pcif1-depleted LLC1 tumours.**

(**a**) Violin plots showing Mki67 expression in single-cell RNA-seq profiles from LLC1 tumours generated from control or Pcif1-depleted cells. Tumours were generated from LLC1 cells expressing a non-targeting shRNA (sh_NTC) or two independent Pcif1 shRNAs (sh_781 and sh_783). Each dot represents one cell. Expression values are shown as scaled z scores. This analysis was used to assess whether Pcif1 depletion was associated with changes in the proliferative state of tumour cells in vivo.

(**b**) Representative multiplex immunofluorescence images of LLC1 tumour sections from the indicated shRNA groups. Sections were stained for CD8a, CD4, F4/80 and CD19 to visualize cytotoxic T cells, helper T cells, macrophages and B cells, respectively. Nuclei are shown in blue. Scale bar, 50 μm. Images are representative of n = 5 tumours per group with similar staining patterns.

(**c**) Representative flow-cytometric gating strategy used to quantify tumour-infiltrating immune populations in LLC1 tumours. Single-cell suspensions were prepared from freshly resected tumours and analysed by flow cytometry. Lymphocyte-enriched events were first selected by forward- and side-scatter properties, followed by exclusion of doublets and dead cells. Live CD45⁺ immune cells were then gated for downstream analysis. T-cell populations were identified within the CD45⁺ compartment, including CD8a⁺ and CD4⁺ T cells, and CD103 and PD-1 expression was assessed within the corresponding T-cell subsets. Myeloid populations were analysed using F4/80, Ly6C and I-A/I-E staining, with F4/80⁺ tumour-associated macrophage populations further resolved by Ly6C and I-A/I-E expression where indicated. The same gating strategy was applied uniformly to all experimental groups.

**Extended Data Figure 6**. **Single-cell analysis of T-cell states in Pcif1-depleted LLC1 tumours.**

(**a**) t-SNE visualization of tumour-infiltrating T cells extracted from the LLC1 single-cell RNA-seq dataset and re-clustered to resolve T-cell states. Annotated subclusters included CD4 T cells, cytotoxic CD8 T cells, progenitor-like exhausted CD8 T cells (CD8Tpex), exhausted CD8 T cells (CD8Tex), γδ T cells, interferon-stimulated gene-positive CD8 T cells (ISG⁺ CD8T), effector-memory CD8 T cells (CD8Tem) and naïve-like CD8 T cells. Each dot represents one cell. Feature plots show expression of Cd8a, Cd4 and Trdc, supporting the identification of CD8 T-cell, CD4 T-cell and γδ T-cell compartments, respectively.

(**b**) Heat map showing z-scored expression of representative genes used to annotate T-cell subclusters. Cytotoxic CD8 T-cell states were characterized by expression of effector molecules and inflammatory mediators, including Gzmb, Ifng, Ccl4, Ccl5, Tnf and Nkg7. ISG⁺ CD8 T cells showed enrichment of interferon-stimulated genes, including Ifit1, Ifit2, Ifit3 and Isg15. Naïve-like or memory-associated T-cell states were marked by Ccr7, Sell, Il7r and related genes. CD4 regulatory features were represented by Foxp3, Ctla4 and Tnfrsf4. γδ T cells were identified by Trdc and Trgc1 expression. Exhaustion-associated CD8 T-cell states were annotated using Tox, Pdcd1, Lag3 and Havcr2. Gene expression values are shown as scaled normalized expression.

(**c**) Stacked bar plot showing the relative abundance of annotated T-cell subclusters among total CD3⁺ T cells in LLC1 tumours generated from shNTC, sh_781 or sh_783 cells. The plot shows the distribution of CD4 T cells, regulatory T cells, cytotoxic CD8 T cells, CD8Tpex, CD8Tex, γδ T cells, ISG⁺ CD8 T cells, CD8Tem and naïve-like CD8 T cells within the tumour-infiltrating T-cell compartment.

(**d**) Violin plots showing expression of selected CD8 T-cell state markers within annotated CD8 T cells from shNTC, sh_781 and sh_783 LLC1 tumours. Tcf7 and Cd69 were used to assess progenitor-like or activation-associated features, whereas Tox, Pdcd1, Entpd1, Havcr2 and Lag3 were used to assess exhaustion-associated features. Each dot represents one CD8 T cell, and violin plots show the distribution of normalized single-cell expression values. P values indicate differences among the three tumour conditions and were calculated using the Kruskal–Wallis test.

(**e**) Violin plots showing expression of selected CD4 T-cell-associated genes within annotated CD4 T cells from shNTC, sh_781 and sh_783 LLC1 tumours. Foxp3 and Ctla4 were used to assess regulatory T-cell-associated features, and [replace with final CD4 marker gene] was used to assess [activation/exhaustion/regulatory/helper-state feature; specify according to the final gene]. Each dot represents one CD4 T cell, and violin plots show the distribution of normalized single-cell expression values. P values indicate differences among the three tumour conditions and were calculated using the Kruskal–Wallis test.

**Extended Data Figure 7**. **PCIF1 loss enhances cholesterol-homeostatic programmes and intracellular cholesterol-associated lipid signals.**

(**a**) Human genetic association analysis linking PCIF1 to metabolic and immune-related traits. Associations between PCIF1 genetic variation and curated human phenotypes were analysed using the Common Metabolic Diseases Knowledge Portal. Traits are grouped by biological category, and each point represents one phenotype. The y axis shows the log-transformed HuGE score, with higher values indicating stronger genetic evidence. Dashed lines indicate reference thresholds for association strength. PCIF1-associated phenotypes included lipid-related, metabolic, cardiovascular, anthropometric, hepatic, renal, immunological and endocrine traits, supporting a broader association between PCIF1 and metabolic homeostasis.

(**b**) Representative polysome-profiling traces from LLC1 and YUMM1.7 cells expressing a non-targeting shRNA (sh_NTC) or two independent Pcif1 shRNAs (sh_781 and sh_783). Cytoplasmic lysates were separated on 5–50% sucrose gradients, and absorbance at 260 nm was monitored across the gradient. The distribution of monosome and polysome fractions was broadly preserved after Pcif1 depletion, indicating that PCIF1 loss did not cause a major collapse of global translation under these conditions. Representative biological replicates are shown for LLC1 and YUMM1.7 cells.

(**c**) Transcriptome-wide comparison of RNA abundance changes in total input RNA and polysome-associated RNA after Pcif1 depletion. The cumulative distribution plot shows log2-transformed fold changes for transcripts in sh_781 cells relative to sh_NTC cells in input and polysome fractions. The box plot summarizes the distribution of log2 fold changes in the two fractions. No significant global shift was detected between input and polysome-associated RNA, indicating that Pcif1 depletion did not broadly remodel transcriptome-wide polysome association. P value was calculated using a two-sided Wilcoxon rank-sum test.

(**d**) Gene set enrichment analysis of RNA-seq data from control and Pcif1-depleted tumour cells showing enrichment of the cholesterol homeostasis gene set in Pcif1-depleted cells. The enrichment plot shows the running enrichment score across the ranked gene list, with normalized enrichment score (NES) and false-discovery rate (FDR q value) indicated. The accompanying heat map shows expression of representative cholesterol-homeostatic and cholesterol-biosynthetic genes, including *Hmgcs1, Sqle, Hmgcr, Hsd17b7, Idi1, Fdft1, Cyp51a1, Stard4, Lss* and *Pmvk*, in sh_NTC, sh_781 and sh_783 cells. Expression values are shown as scaled z-score normalized expression.

(**e**) Schematic of the cholesterol-biosynthesis pathway highlighting genes increased after Pcif1 depletion. Genes shown in red were upregulated in Pcif1-depleted cells and validated by both RNA-seq and quantitative PCR analysis. The highlighted enzymes span the mevalonate pathway, isoprenoid synthesis, squalene synthesis and downstream sterol-biosynthesis steps, supporting coordinated induction of the SREBP2-associated cholesterol-biosynthetic programme after PCIF1 loss. This schematic summarizes transcriptional activation of cholesterol-biosynthetic genes and does not indicate direct measurement of metabolic flux.

(**f**) Flow-cytometric gating strategy for Filipin III analysis of free cholesterol in tumour cells. Tumour cells were first selected by forward- and side-scatter properties, followed by gating of single cells and live cells. Filipin III fluorescence was then quantified in the live single-cell population. Representative histograms show negative-control and Filipin III-positive populations used to define cholesterol-associated fluorescence. The same gating strategy and acquisition settings were applied across all experimental groups.

(**g**) Representative confocal images of LLC1 cells expressing sh_NTC, sh_783 or sh_781 stained with Filipin III, BODIPY C16 and PCIF1. Filipin III was used to detect free cholesterol, BODIPY C16 was used to visualize lipid-associated compartments and assess their spatial relationship with Filipin III-positive signals, and PCIF1 staining confirmed reduced PCIF1 signal in Pcif1-depleted cells. Overlay images show increased Filipin III signal in Pcif1-depleted cells, with partial spatial overlap between Filipin III and BODIPY C16 signals. Scale bars, 10 μm. Images are representative of n = 3 independent experiments with similar results.

**Extended Data Figure 8**. **Lipidomic profiling of Pcif1-depleted LLC1 cells.**

(**a**) Experimental workflow for untargeted lipidomic profiling of LLC1 cells. Control LLC1 cells and Pcif1-depleted LLC1 cells were collected under matched culture conditions and subjected to liquid chromatography–mass spectrometry-based lipidomics. Lipid features were detected according to mass-to-charge ratio (m/z), retention time and ion intensity and analysed separately in negative- and positive-ion acquisition modes.

(**b**) Representative total ion chromatograms from the lipidomics analysis in negative- and positive-ion modes. Chromatograms are shown for solvent blank (N/A), pooled quality-control (QC) sample and representative biological sample. The solvent blank was used to monitor background and carryover signals, whereas pooled QC samples were used to evaluate chromatographic stability and feature reproducibility during acquisition. Retention time is shown on the x axis, and total ion intensity is shown on the y axis.

(**c**) Density distribution of detected lipid-feature intensities across the lipidomics dataset before downstream differential analysis. The distribution was used to assess the overall signal range and intensity structure of detected lipid features.

(**d**) Density distribution of normalized lipid-feature intensities after data processing and normalization. The normalized intensity distribution was used to assess comparability of lipid-feature abundance across biological samples before multivariate and differential lipidomic analyses.

(**e**) Sparse partial least-squares discriminant analysis (sPLS-DA) of lipidomic profiles from Pcif1-depleted and control LLC1 cells. Negative-ion mode data are shown at the top, and positive-ion mode data are shown at the bottom. Each point represents one biological sample. Shaded ellipses indicate group distribution. Separation between Pcif1-depleted and control cells indicates that PCIF1 loss is associated with broad remodelling of the cellular lipidome in both ionization modes.

(**f**) Differential lipid-feature analysis in negative-ion mode comparing Pcif1-depleted LLC1 cells with control cells. The volcano plot shows log2-transformed fold change on the x axis and −log10-transformed P value on the y axis. Lipid features were considered significantly altered when they met both thresholds: FDR < 0.05 and |log2 fold change| > 1.5. Features increased in Pcif1-depleted cells are shown in red, features decreased in Pcif1-depleted cells are shown in green, and non-significant features are shown in grey. The heat map shows normalized intensities of significantly altered lipid features across individual samples, with unsupervised clustering of samples and lipid features.

(**g**) Differential lipid-feature analysis in positive-ion mode comparing Pcif1-depleted LLC1 cells with control cells. The volcano plot shows log2-transformed fold change and −log10-transformed P value for each detected lipid feature. Significantly altered lipid features were defined using FDR < 0.05 and |log2 fold change| > 1.5. Increased, decreased and non-significant lipid features are shown as in f. The accompanying heat map displays normalized intensities of significantly altered features across biological samples, showing lipidomic separation between Pcif1-depleted and control LLC1 cells.

(**h**) Lipid-class-level summary of differential lipid features detected in negative-ion mode. The bubble plot shows the median log2 fold change for each lipid class in Pcif1-depleted cells relative to control cells on the x axis and the class-level false-discovery rate on the y axis. Class-level significance was calculated by aggregating feature-level P values within each lipid class, followed by false-discovery-rate correction. Bubble size indicates the number of detected features in each lipid class. The bar plot shows the number of significantly downregulated and upregulated features within each lipid class. Lipid classes detected in negative-ion mode included phosphatidylethanolamine (PE), phosphatidylglycerol (PG), phosphatidylcholine (PC), phosphatidylinositol (PI), cardiolipin (CL), sphingomyelin (SM), bis(monoacylglycero)phosphate/hemibismonoacylglycerophosphate (HBMP), glucosylceramide (GlcCer), phosphatidic acid (PA), ceramide (Cer), phosphatidylserine (PS) and fatty acids (FA).

(**i**) Lipid-class-level summary of differential lipid features detected in positive-ion mode. The bubble plot shows the median log2 fold change and class-level false-discovery rate for each lipid class, and the bar plot shows the number of downregulated and upregulated features in Pcif1-depleted cells relative to control cells. Class-level significance was calculated by aggregating feature-level P values within each lipid class, followed by false-discovery-rate correction. Lipid classes detected in positive-ion mode included phosphatidylcholine (PC), triacylglycerol (TAG), phosphatidylethanolamine (PE), sphingomyelin (SM), glucosylceramide (GlcCer), phosphatidylserine (PS), acylcarnitine (ACar) and phosphatidylglycerol (PG). Together, these analyses show that Pcif1 depletion is accompanied by broad lipidomic remodelling, including changes in phospholipid, sphingolipid, glycerolipid and mitochondrial lipid-associated classes. Class-level significance was calculated by combining feature-level P values within each lipid class using Fisher’s combined probability test, followed by Benjamini-Hochberg false-discovery-rate correction across lipid classes.

**Extended Data Figure 9**. **PCIF1 loss enhances SCAP translational engagement and links cholesterol sensing to NFκB-associated inflammatory signalling.**

(**a**) Quantitative PCR analysis of genes involved in SCAP–SREBP2 cholesterol sensing, ER-to-Golgi transport and SREBP2 proteolytic processing in Pcif1-depleted tumour cells. Left, schematic of SCAP-dependent SREBP2 activation. Under cholesterol-permissive conditions, SCAP escorts SREBP2 from the endoplasmic reticulum to the Golgi, where SREBP2 is sequentially cleaved by site-1 and site-2 proteases to release the N-terminal transcription factor domain. The cleaved SREBP2-N fragment translocates to the nucleus and induces cholesterol-biosynthetic gene expression. Right, qPCR analysis of Scap, Sec24a, Insig2, Mbtps1, Mbtps2 and Sar1a mRNA expression in YUMM1.7 and LLC1 cells expressing a non-targeting shRNA (shNTC) or two independent Pcif1 shRNAs (sh_781 and sh_783). Expression values were normalized to [housekeeping gene] and are shown as mean ± s.e.m.; each dot represents one biologically independent sample. P values were calculated using unpaired Mann-Whitney test. ns, not significant. These data indicate that PCIF1 loss does not uniformly increase mRNA abundance of SCAP-associated trafficking or SREBP2-processing components, supporting the interpretation that increased SCAP protein abundance and SREBP2 activation are not simply explained by broad transcriptional induction of this machinery.

(**b**) xTAIL analysis of polysome RNA-seq data from LLC1 and YUMM1.7 cells after Pcif1 depletion. Total input RNA and polysome-associated RNA were profiled by RNA-seq from shNTC, sh_781 and sh_783 cells, and changes in translational efficiency were inferred using xTAIL. The panel will show [final plot type: for example, volcano plots, scatter plots or ranked translational-efficiency plots] for LLC1 and YUMM1.7 cells, highlighting transcripts with significant changes in translational efficiency after Pcif1 depletion. Significant translational-efficiency changes were defined using [FDR threshold] and [effect-size cutoff]. Scap is indicated in the corresponding analysis where applicable. This analysis was used to determine whether PCIF1 loss selectively alters translational engagement of specific transcripts, rather than inducing a broad collapse of polysome-associated translation.

(**c**) RNA-seq quantification of Scap abundance in total input RNA and polysome-associated RNA from LLC1 and YUMM1.7 cells expressing shNTC, sh_781 or sh_783. Bar plots show Scap counts per million (CPM) in total RNA and polysome-associated RNA fractions. Pcif1 depletion increased Scap abundance in the polysome-associated fraction, whereas Scap abundance in total RNA was not increased to the same extent, supporting enhanced translational engagement of Scap after PCIF1 loss. Data are shown as mean ± s.e.m.; each dot represents one biological replicate. P values for the indicated comparisons were calculated using one-way ANOVA test.

(**d**) Immunoblot analysis of SCAP protein abundance and SREBP2 processing in YUMM1.7 cells expressing shNTC, sh_781 or sh_783. Precursor SREBP2 (SREBP2-P), cleaved N-terminal SREBP2 (SREBP2-N), PCIF1 and SCAP were detected by immunoblotting. α-actin and β-tubulin were used as loading controls for the corresponding blots. Increased SCAP protein and SREBP2-N abundance in Pcif1-depleted cells support activation of the SCAP–SREBP2 cholesterol-biosynthetic axis after PCIF1 loss. Blots are representative of n = 3 independent experiments with similar results.

(**e**) Distribution of the control transcript *Hprt1* across polysome-gradient fractions in YUMM1.7 cells. Cytoplasmic lysates from shNTC, sh_781 and sh_783 cells were separated on 5–50% sucrose gradients, and Hprt1 mRNA abundance in each fraction was quantified by qPCR. Values are expressed as the percentage of total Hprt1 mRNA recovered across all fractions. Hprt1 showed broadly similar distribution across polysome fractions in control and Pcif1-depleted cells, indicating that PCIF1 loss did not induce a general shift in the polysome association of this control transcript.

(**f**) Representative polysome-profiling trace from LLC1 cells expressing shNTC, sh_781 or sh_783. Cytoplasmic lysates were separated on a 5–50% sucrose gradient, and absorbance at 260 nm was recorded across the gradient. The overall distribution of monosome and polysome fractions was broadly preserved after Pcif1 depletion, consistent with the absence of a major global collapse of translation under these conditions.

(**g**) Distribution of *Scap* mRNA across polysome-gradient fractions in LLC1 cells. After sucrose-gradient fractionation, Scap mRNA abundance in each fraction was quantified by qPCR and expressed as the percentage of total Scap mRNA recovered across all fractions. Compared with shNTC cells, Pcif1-depleted cells showed increased representation of Scap mRNA in heavier polysome fractions, supporting enhanced polysome association of Scap after PCIF1 loss.

(**h**) Distribution of *Hprt1* mRNA across polysome-gradient fractions in LLC1 cells analysed as in g. Hprt1 was used as a control transcript to assess whether Pcif1 depletion broadly altered mRNA distribution across the sucrose gradient. The Hprt1 distribution was not shifted to the same extent as Scap, supporting transcript-selective enhancement of Scap polysome association.

(**i**) Immunoblot analysis of NFκB p65 phosphorylation in YUMM1.7 control and Pcif1-depleted cells treated with vehicle or UT-59. UT-59 was used to inhibit SCAP-dependent SREBP2 activation. Phospho-p65, total p65 and PCIF1 were detected by immunoblotting, with β-tubulin used as a loading control. UT-59 attenuated the increase in p65 phosphorylation associated with PCIF1 loss, supporting a role for SCAP-dependent cholesterol sensing upstream of NFκB-associated inflammatory signalling. Blots are representative of n = 2 independent experiments with similar results.

(**j**) Immunoblot analysis of NFκB p65 phosphorylation in LLC1 control and Pcif1-depleted cells treated with vehicle or methyl-β-cyclodextrin (MβCD). MβCD was used to reduce accessible cellular cholesterol. Phospho-p65, total p65 and PCIF1 were detected by immunoblotting, with β-tubulin used as a loading control. MβCD reduced the PCIF1-loss-associated increase in p65 phosphorylation, indicating that cholesterol availability contributes to NFκB activation in Pcif1-depleted tumour cells. Blots are representative of n = 2 independent experiments with similar results.

(**k**) Immunoblot analysis of NFκB p65 phosphorylation in YUMM1.7 and LLC1 control and Pcif1-depleted cells treated with vehicle or U18666A. U18666A blocks NPC1-dependent cholesterol egress from late endosomes and lysosomes and was used as an orthogonal perturbation of intracellular cholesterol handling. Phospho-p65, total p65 and PCIF1 were detected by immunoblotting, with β-tubulin used as a loading control. U18666A altered p65 phosphorylation under the indicated conditions, supporting a link between intracellular cholesterol handling and NFκB-associated inflammatory signalling. Blots are representative of n = 2 independent experiments with similar results.

(**l**) Standard curve for CXCL10 ELISA quantification. Recombinant CXCL10 standards were serially diluted across the indicated concentration range, and absorbance was measured at 450 nm with 630 nm background subtraction. The standard curve was fitted using a four-parameter logistic regression model. Best-fit parameters are shown in the panel and were used to interpolate CXCL10 concentrations in conditioned-medium samples.

**Extended Data Figure 10**. **Clinical association of PCIF1 expression with checkpoint response, cytolytic immune activity and cancer outcome, and proposed model of the PCIF1–SCAP–cholesterol inflammatory axis.**

(**a**) Kaplan–Meier analyses of overall survival and progression-free survival in melanoma patients treated with anti-PD1 therapy, stratified by tumour PCIF1 expression. Patients were divided into PCIF1-low and PCIF1-high groups using the optimal expression cutoff selected by the survival-analysis platform. Hazard ratios (HRs), 95% confidence intervals and log-rank P values are shown. Numbers at risk are indicated below each plot. High PCIF1 expression was associated with shorter overall survival and progression-free survival in anti-PD1-treated melanoma patients.

(**b**) Kaplan–Meier analyses of overall survival and progression-free survival in melanoma patients treated with anti-CTLA4 therapy, stratified by tumour PCIF1 expression as in a. HRs, 95% confidence intervals, log-rank P values and numbers at risk are shown. High PCIF1 expression was associated with reduced survival outcomes in this independent checkpoint-blockade context.

(**c**) Kaplan–Meier analysis of overall survival in glioblastoma patients treated with anti-PD1 therapy, stratified by PCIF1 expression. Patients were divided into PCIF1-low and PCIF1-high groups using the optimal expression cutoff. HR, 95% confidence interval, log-rank P value and numbers at risk are shown. High PCIF1 expression was associated with shorter overall survival.

(**d**) Kaplan–Meier analysis of progression-free survival in glioblastoma patients treated with anti-PD1 therapy, stratified by PCIF1 expression as in c. HR, 95% confidence interval, log-rank P value and numbers at risk are shown. High PCIF1 expression was associated with shorter progression-free survival.

(**e**) Association between PCIF1 expression and cytolytic immune activity across tumour cohorts. Scatter plots show PCIF1 expression versus a gene-expression-derived cytolytic T-cell activity score (CTL) in glioblastoma from TCGA, lung adenocarcinoma from GSE30219, breast cancer from METABRIC and stage IV melanoma from GSE22153. Each dot represents one tumour sample. Dashed lines indicate linear regression fits. Across these cohorts, higher PCIF1 expression was associated with lower CTL scores, consistent with an inverse relationship between tumour PCIF1 expression and cytolytic immune activity.

(**f**) Kaplan–Meier analyses of overall survival in lung adenocarcinoma (LUAD) and colon adenocarcinoma (COAD) patients stratified by PCIF1 expression. Patients were divided into PCIF1-low and PCIF1-high groups using the optimal expression cutoff. HRs, 95% confidence intervals, log-rank P values and numbers at risk are shown. High PCIF1 expression was associated with shorter overall survival in both cohorts.

(**g**) Kaplan–Meier analyses of overall survival in oestrogen receptor-positive breast cancer (ER⁺ BRCA) and serous ovarian cancer patients stratified by PCIF1 expression. Patients were divided into PCIF1-low and PCIF1-high groups using the optimal expression cutoff. HRs, 95% confidence intervals, log-rank P values and numbers at risk are shown. High PCIF1 expression was associated with shorter overall survival in both tumour types.

(**h**) Proposed model of tumour-cell PCIF1 function in cholesterol-linked antitumour immunity. In PCIF1-intact tumour cells, SCAP translational engagement and SCAP–SREBP2 pathway activity are restrained, resulting in limited cholesterol-biosynthetic gene expression and low intracellular cholesterol accumulation. This state is associated with an immune-cold tumour microenvironment, low CD8⁺ T-cell infiltration and poor response to immune checkpoint blockade. Upon PCIF1 loss, SCAP translational engagement increases together with SREBP2 activation, cholesterol-biosynthetic gene induction and intracellular cholesterol accumulation. This cholesterol-associated state is accompanied by an inflammatory cytokine programme, including CXCL10, which promotes CD8⁺ T-cell infiltration, tumour control and improved checkpoint responsiveness. Clinically, high PCIF1 expression is associated with poorer ICB outcome, whereas low PCIF1 expression or PCIF1 loss is associated with higher cytolytic immune activity.
